# LIN-44/Wnt controls developmental neurite pruning via UNC-43/CaMKII and PKC-2/PKC in *C. elegans*

**DOI:** 10.64898/2026.06.04.729963

**Authors:** Menghao Lu, Jeffrey S.H. Lin, Mizuki Kurashina, Kota Mizumoto

**Affiliations:** Department of Zoology; Graduate program in Cell and Developmental Biology; Life Sciences Institute; Djavad Mowafaghian Centre for Brain Health, The University of British Columbia, Vancouver, Canada, V6T 1Z3

**Keywords:** Neurite pruning, Wnt, IP3R, Calcium signaling, CaMKII, PKC, Endocytosis, *C*. *elegans*

## Abstract

During development, many neurons prune their neurites. While many pruning events are activity-dependent, some neurons undergo stereotyped and developmentally regulated neurite pruning, and our understanding of the signaling pathways that mediate this form of pruning remains limited. In this study, using the PDB motor neuron in *C. elegans*, we show that the Wnt-calcium signaling pathway is required for stereotyped neurite pruning during development. We found that mutants of *itr-1/*IP3 receptor and two calcium-dependent kinases, *unc-43/*CaMKII and *pkc-2/*PKC, exhibit neurite pruning defects. Genetic analysis suggested that they function downstream of *lin-44/*Wnt in neurite pruning. Human CaMKIIA can induce neurite pruning in *C. elegans*, and mutations in CaMKII genes in patients with intellectual disabilities affect its pruning function. In vivo calcium imaging revealed that PDB neurites exhibit calcium transients during neurite pruning, which are regulated at least in part by *lin-44* and *itr-1*. Furthermore, we demonstrate that *pkc-2* regulates neurite pruning through clathrin-mediated endocytosis. Together, our work reveals the critical functions of Wnt–calcium signaling in neurite pruning.

## Introduction

Developmental neurite pruning is an essential process for refining the nervous system. During development, neurons extend excessive neurites, many of which are later severed or retracted (Luo & O’Leary, 2005; Riccomagno & Kolodkin, 2015; Schuldiner & Yaron, 2015). Neurite pruning can occur in an activity-dependent manner, in which neurites with weaker synaptic input/output are selectively eliminated. For instance, monocular deprivation in four-week-old mice induces axon and dendritic spine pruning in the visual cortex corresponding to the deprived eye, which is mediated by the inhibitory neurotransmitter γ-aminobutyric acid (GABA) (Antonini & Stryker, 1993; Hensch, 2005; Yamahachi et al., 2009; Zhou et al., 2017). Similarly, in *Drosophila*, the GABA-mediated reduction of neuronal activity in mushroom γ-Kenyon cells is required for its axonal pruning (Mayseless et al., 2023). On the other hand, some developmental cues initiate more stereotyped neurite pruning. For example, during the metamorphosis of *Drosophila*, ecdysone signaling induces dendrite pruning in class IV da neurons (Kuo et al., 2005), axon pruning in the mushroom body APL neuron (Mayseless et al., 2018), and neurite pruning in Beat-VaM neurons (Lehmann et al., 2025). In the mouse hippocampus, stereotyped pruning of the mossy fiber axons of the infrapyramidal bundle occurs between postnatal day 10 and day 45, and is regulated by Semaphorin (Bagri et al., 2003) and Ephrin (Xu & Henkemeyer, 2009) signaling pathways. Despite these findings, our understanding of stereotyped developmental neurite pruning remains limited, possibly due to technical difficulties in observing pruning events in live animals.

The simple yet highly conserved nervous system of *Caenorhabditis elegans* serves as an advantageous model for dissecting the genetic mechanisms underlying developmental neurite pruning, owing to its stereotyped development and transparent body, which allow direct observation of the process in live animals. Several neurons in *C. elegans* have been shown to exhibit neurite pruning. For example, the AIM interneuron exhibits stereotyped neurite pruning, a process which is promoted by the transcription factor MBR-1/LCOR (Kage et al., 2005), and is inhibited by two Wnt morphogens, CWN-1 and CWN-2 (Hayashi et al., 2009). We previously reported that the developmental neurite pruning of a post-embryonic cholinergic motor neuron, PDB, is regulated by LIN-44/Wnt and its receptor, LIN-17/Frizzled (Lu & Mizumoto, 2019). During its development, the PDB neuron initially extends transient neurites posteriorly towards the tail, where LIN-44/Wnt-expressing hypodermal cells reside. Once posteriorly growing neurites are in close proximity to the LIN-44/Wnt-expressing cells, they undergo stereotyped pruning during the second (L2) to the third larval stages (L3) via retraction. Interestingly, membrane-tethered LIN-44 is sufficient to induce neurite pruning of PDB, suggesting the diffusion and gradient distribution of LIN-44 is dispensable to induce neurite pruning (Lu & Mizumoto, 2019).

Wnt signaling plays crucial roles in neurodevelopment, including neurogenesis, neuronal polarization, axon guidance, synapse formation, and synaptic plasticity (He et al., 2018; Park & Shen, 2012; Salinas & Zou, 2008). Upon Wnt binding, Frizzled receptors activate three distinct downstream signaling cascades: canonical β-catenin signaling, planar polarity pathway (PCP) signaling, and Wnt-calcium signaling (He et al., 2018). Among them, the Wnt-calcium pathway remains less understood, especially in the nervous system. Wnt-calcium signaling was first observed in the zebrafish embryo, where Wnt-5a overexpression stimulates intracellular calcium signaling in the superficial cells of the blastodisc through its receptor Frizzled-2, thereby inducing cell movement and ventralization (Slusarski, Yang-Snyder, et al., 1997; Westfall et al., 2003). The Wnt-calcium pathway involves activation of phospholipase C (PLC) (Sheldahl et al., 2003; Slusarski, Corces, et al., 1997). Active PLC facilitates the production of inositol 1,4,5-triphosphate (IP3) to elevate intracellular calcium level, which then activates downstream components such as calcium/calmodulin-dependent protein kinase II (CaMKII), protein kinase C (PKC), and calcineurin (Heisenberg et al., 2000; Kuhl, Sheldahl, Malbon, et al., 2000; Saneyoshi et al., 2002; Sheldahl et al., 1999; Sheldahl et al., 2003). In *C.elegans*, two Wnts, LIN-44 and EGL-20, are found to regulate anal depressor muscle remodeling by activating calcium signal via IP3 signaling and ryanodine receptors (RyR) through *egl-8/*PLCβ (LeBoeuf et al., 2020).

A few *in vitro* studies suggest a role of the Wnt-calcium pathway in regulating axon outgrowth and synapse development. For example, Wnt5a treatment to the dissociated neurons from the hamster sensorimotor cortex promotes axon outgrowth and repels growth cones in an IP3- and CaMKII-dependent manner (Hutchins et al., 2011; Li et al., 2009). Wnt5a treatment also increases the dendritic calcium level in dissociated rat hippocampal neurons through voltage-gated calcium channel (VGCCs) and regulates GluN2B-containing synaptic N-methyl-D-aspartate receptor (NMDAR) trafficking (McQuate et al., 2017). However, the roles of the Wnt-calcium pathway in neurite pruning remain elusive.

In this study, we report that the Wnt-calcium pathway regulates developmental neurite pruning in *C. elegans*. Through genetic analysis, we describe that LIN-44/Wnt acts through ITR-1/inositol 1,4,5-triphosphate receptor, UNC-43/CaMKII and PKC-2/PKC. In addition, we find that PKC-2 regulates neurite pruning through clathrin-mediated endocytosis. Taken together, this study provides a novel function of the Wnt-calcium pathway in regulating developmental neurite pruning.

## Results

### *unc-43/*CaMKII functions downstream of *lin-44/*Wnt to regulate PDB neurite pruning

The PDB neuron extends a single neurite posteriorly along the ventral side of the animal and makes a characteristic ‘V-shape’ turn to join the dorsal nerve cord, where it forms *en passant* neuromuscular junctions with dorsal body wall muscles (White et al., 1986). Previously, we showed that the turning of the PDB neurite involves neurite pruning (Lu & Mizumoto, 2019). During the second larval (L2) stage, PDB forms transient neurites extending posteriorly at the turning point, most of which are pruned via retraction by the last larval stage (L4) (**Figure 1A**) (Lu & Mizumoto, 2019). In the *lin-44(n1792)* null mutant, the posteriorly extending neurites often fail to prune, resulting in the presence of posterior neurites at the L4 stage (**Figure 1B**) (Lu & Mizumoto, 2019).

**Figure 1.**
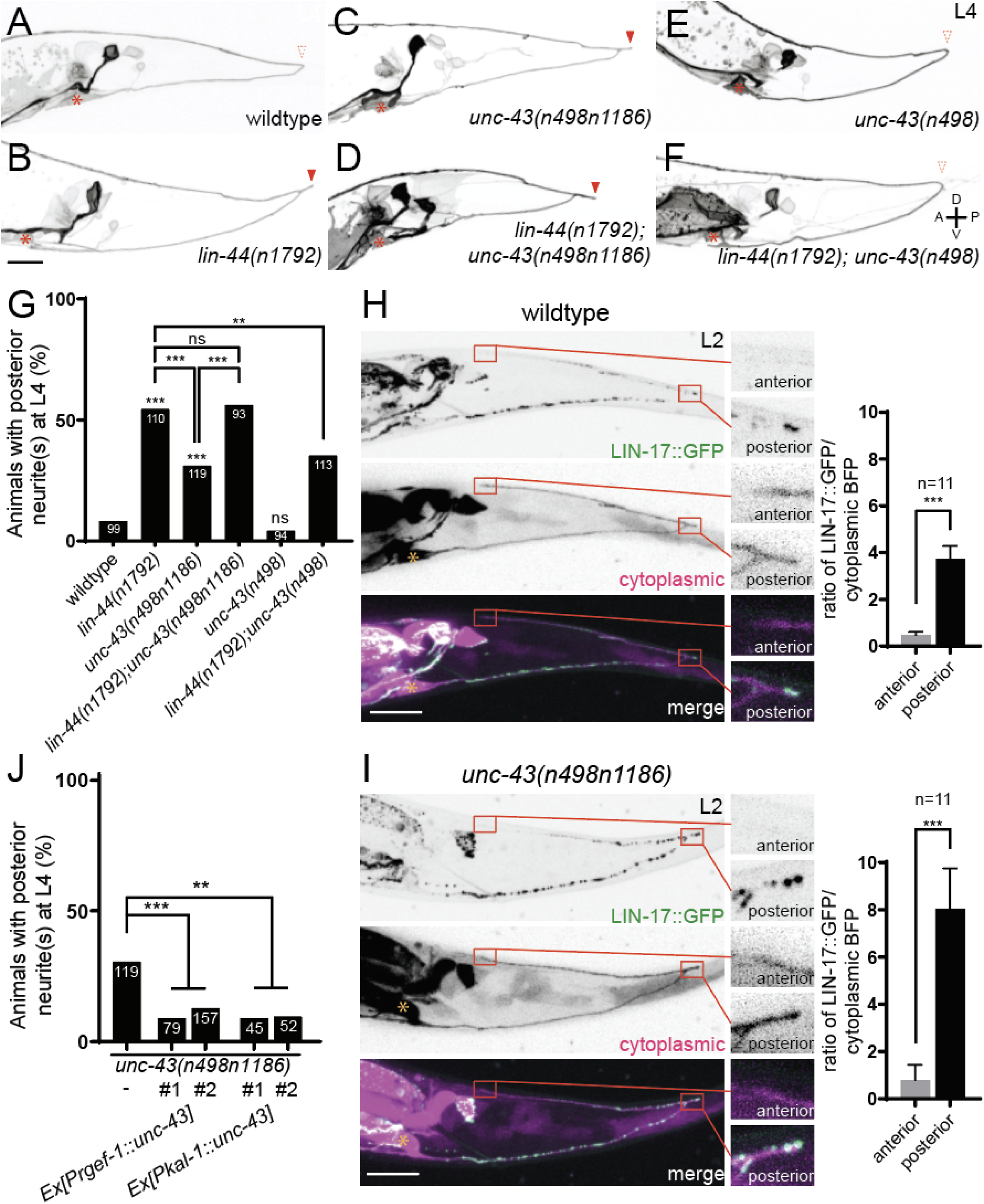
*unc-43* functions downstream of *lin-44* in proper PDB development. **(A-F)** Representative images of PDB at the L4 stage. Red arrowheads denote posterior neurites, and dotted arrowheads represent the absence of posterior neurites. Asterisks denote the PDB cell bodies. **(G)** Quantification of animals with ectopic posterior neurites. Sample numbers are shown in each bar. ***p < 0.001; **p < 0.002; ns, not significant (Fisher’s exact test). **(H, I)** Representative images of LIN-17::GFP localization (top panels), PDB neurite structure labeled with cytoplasmic BFP (middle panels) and merged images (bottom panels) in wildtype **(H)** and *unc-43(n498n1186)* **(I)** animals. Quantification of the normalized GFP/BFP ratio at the anterior and posterior growth cones for each genotype is shown on the right. Error bars indicate mean ± SEM. ***p < 0.001 (Ratio paired t-test). **(J)** Quantification of the animals with ectopic posterior neurites. Two independent lines of each rescue experiment are labeled with #1 and #2. Note the quantification of *unc-43(n498n1186)* is from panel (**G**). Sample numbers are shown in each bar. ***p < 0.001; *p < 0.033 (Fisher’s e2xact test). Scale bars: 10 μm.

To identify the mechanisms of Wnt-dependent neurite pruning, we screened for mutants with posterior neurites at the L4 stage. In wildtype, 8% (8/99) of animals have ectopic posterior branches in PDB, while 55% (60/110) of *lin-44(n1792)* mutant animals have ectopic branches (**Figure 1G**). Our previous screening suggested neither β-catenin nor the Planar Cell Polarity (PCP) pathway acts downstream of LIN-44/Wnt (Lu & Mizumoto, 2019). This prompted us to examine the components of the Wnt-calcium signaling pathway. In Xenopus, non-canonical Wnts, Xwnt-5A and Xwnt-11, activate CaMKII for ventral fate specification during embryogenesis (Kuhl, Sheldahl, Malbon, et al., 2000). Consistently, we found that 31% of null mutants of *unc-43*, the sole ortholog of CaMKII in *C. elegans* (Reiner et al., 1999), had posterior neurites in PDB at the L4 stage (n =119) (**Figures 1C and 1G**). To determine whether the posterior neurites observed in *unc-43(n498n1186)* mutants were due to the pruning defects, we examined the neurite pruning frequency. We selected animals with posterior neurites at the L2 stage and re-examined their PDB morphology at the L4 stage, as we did previously (Lu & Mizumoto, 2019). We found that 46% (n = 61) of the posterior neurites observed at the L2 stage did not prune in *unc-43(n498n1186)* mutants, which is significantly higher than the neurite pruning frequency in wildtype (14%, n = 63) (**Figures 2A, 2B, 2E, 2F**). This result indicates that the posterior neurites observed at the L4 stage of *unc-43(n498n1186)* mutant animals are caused by the pruning defect.

**Figure 2.**
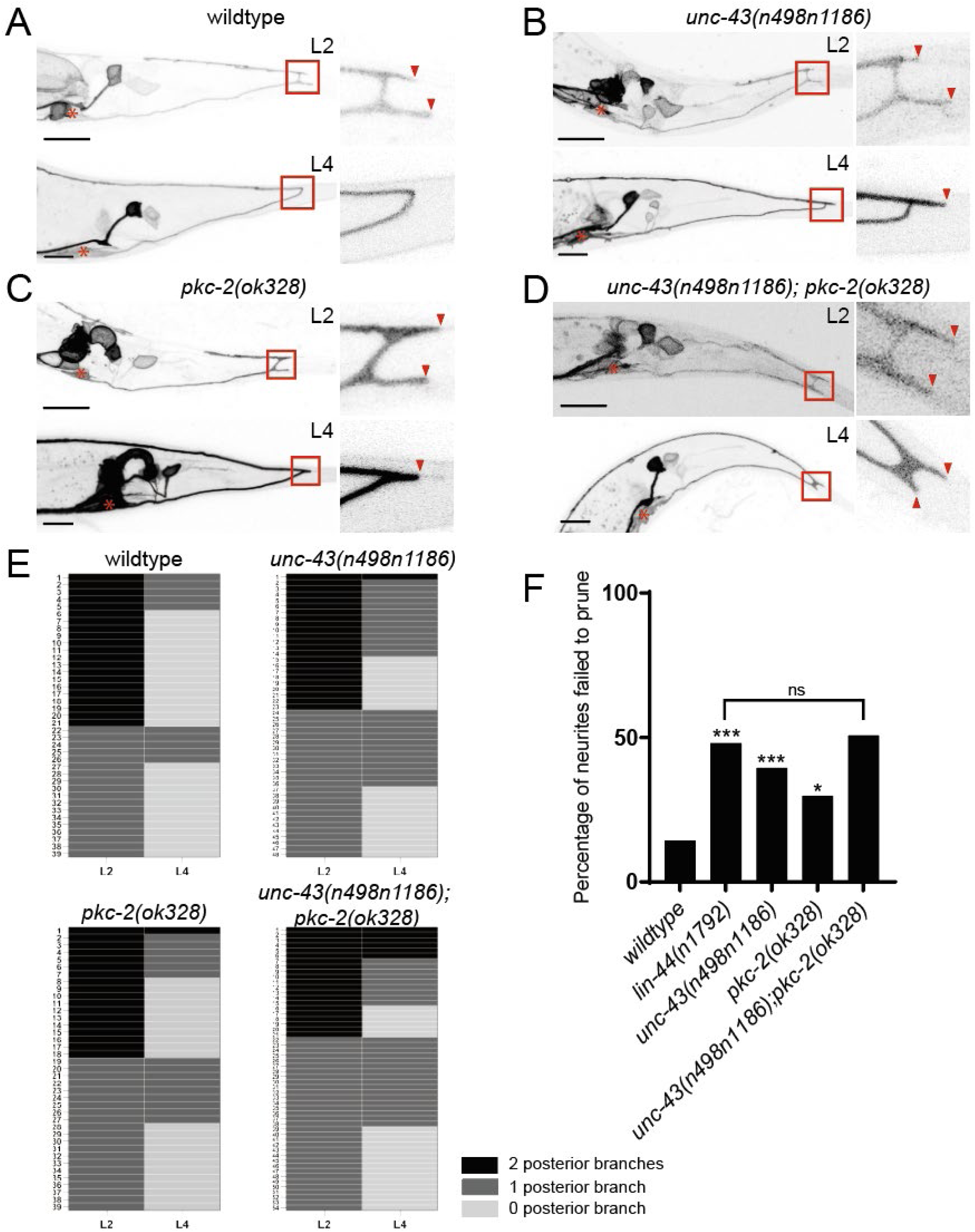
*unc-43* and *pkc-2* are required for neurite pruning of PDB. **(A-D)** Representative images of single animals of wildtype **(A)**, *unc-43(n498n1186)* **(B)**, *pkc-2(ok328)* **(C)**, and *unc-43(n498n1186); pkc-2(ok328)* double mutant **(D)** at the L2 (top panels) and the L4 stages (bottom panels). Red arrowheads denote the posterior neurites. Asterisks denote the PDB cell bodies. **(E)** Quantification of the posterior neurite number of individual animals at L2 and L4 stages in each genetic background. **(F)** Quantification of the posterior neurite pruning frequency. ***p < 0.001; *p < 0.033; ns, not significant (Fisher’s exact test). Scale bars: 10 μm.

To determine the relationship between *unc-43/CaMKII* and *lin-44/Wnt* in neurite pruning, we examined their genetic epistasis. The penetrance of posterior neurite at L4 in the *lin-44; unc-43* double mutants (56%, n=93) was not significantly different from that in *lin-44* single mutant (55%, n=110), suggesting that *unc-43* functions in the same genetic pathway as *lin-44* (**Figures 1D, 1G**). The gain-of-function mutant of *unc-43(n498)*, which carries an E108K mutation causing Ca^2+^ -independent activation of UNC-43 (Robatzek & Thomas, 2000), does not exhibit posterior neurite phenotype at the L4 stage (4%, n = 94) **(Figures 1E, 1G)**. In addition, *unc-43(n498)* partially suppressed the *lin-44(n1791)* mutant, suggesting that *unc-43* acts downstream of *lin-44* (**Figures 1F, 1G**). We previously reported that LIN-17/Frizzled is localized at the posterior neurites during the L2-L3 stages (**Figure 1H**) in a LIN-44-dependent manner (Lu & Mizumoto, 2019). In *unc-43(n498n1186)* null mutants, LIN-17 localization pattern is unaffected, which is consistent with the idea that *unc-43* acts downstream of *lin-44/Wnt* and *lin-17/Frizzled* (**Figures 1H, 1I**). Expression of *unc-43* cDNA under the pan-neuronal promoter (P*rgef-1*) or the PDB-specific promoter (P*kal-1*) rescued the posterior neurite phenotype observed at L4 in *unc-43(n498n1186)* mutants, suggesting that *unc-43* functions cell-autonomously in PDB to regulate neurite pruning **(Figure 1J)**. Taken together, we conclude that *unc-43* functions in PDB and acts downstream of *lin-44/Wnt* and *lin-17/Frizzled* in neurite pruning.

### UNC-43/CaMKII is enriched at the pruning neurites of PDB in a Wnt-independent manner

We next examined the subcellular localization of endogenous UNC-43 in PDB. We took advantage of the NATF (Native and Tissue-Specific Fluorescence) system (He et al., 2019) by knocking in seven tandem repeats of *GFP_11_* sequence at the 5’ end of the *unc-43* locus using CRISPR/Cas9. By expressing GFP_1-10_ in PDB using the PDB-specific promoter (P*kal-1*), the fluorescent GFP is reconstituted to visualize endogenous UNC-43 protein in PDB. We found that 7×GFP_::_UNC-43 puncta are enriched in the posterior neuronal processes of PDB, including the posterior neurites, during the L2-L3 stages when the pruning occurs **(Figure S1A**). Interestingly, the 7×GFP_::_UNC-43 signal is diminished at the L4 stage or in adult animals when the pruning of PDB has completed **(Figure S1A)**. The enrichment of 7×GFP_::_UNC-43 at the posterior neurites in L2-L3 stage animals is unaffected in the *lin-44(n1792)* mutant (**Figure S1B**), suggesting that LIN-44 does not regulate neurite pruning via UNC-43 localization but possibly by other mechanisms, such as regulating the kinase activity of UNC-43.

### Human CaMKIIA can functionally replace *unc-43* in PDB neurite pruning

Previous work demonstrated that transgenic expression of human CaMKIIA cDNA can rescue the diffuse synaptic vesicle localization phenotype of the *unc-43(e408)* mutant (Chia et al., 2018). To test whether human CaMKIIA can also replace the function of *unc-43* in neurite pruning, we generated a *C. elegans* strain in which the entire coding region of *unc-43* was replaced with the codon-optimized human CaMKIIA (hCaMKIIA) sequence using CRISPR/Cas9. The resulting *unc-43(miz214[hCaMKIIA])* animals had normal PDB structure with no significant posterior neurite phenotype, suggesting human CaMKIIA can induce neurite pruning similar to UNC-43 (**Figures S2A, S2E**).

Previous work has identified a H477Y loss-of-function mutation in the CaMKIIA in patients with a severe form of intellectual disability (Chia et al., 2018). The *unc-43(miz214miz238[hCaMKIIA H477Y])* mutant exhibits the posterior neurite phenotype similar to that of *unc-43(n498n1186)* null mutants, suggesting that the H477Y mutation abolishes the function of CaMKIIA in neurite pruning (**Figures S2B, S2E**). Conversely, the *unc-43(miz214miz239[hCaMKIIA K291P])* mutant carrying a gain-of-function mutation (K291P), which is equivalent to the R292P mutation found in CaMKIIG of patients with intellectual disability (Proietti Onori et al., 2018), suppressed the posterior neurite phenotype of *lin-44(n1792)* mutant (**Figure S2C-S2E**). Taken together, these findings suggest that human CaMKIIA may have a conserved function in regulating neurite pruning.

### *pkc-2/*PKC functions in parallel to *unc-43/*CaMKII to regulate PDB neurite pruning

The penetrance of the posterior neurite phenotype in *unc-43(n498n1186)* mutants is lower than in *lin-44(n1792)* mutants (**Figure 1G**). This raises the possibility that additional factor(s) may function in parallel to *unc-43* to regulate PDB neurite pruning. Protein kinase C (PKC) is another calcium-dependent kinase known to be regulated by Wnt signaling (Kuhl, Sheldahl, Park, et al., 2000). Consistently, the null mutant of *pkc-2(ok328)/PKC* exhibits a mild but significant posterior neurite phenotype at the L4 stage (21%, n=101), while *pkc-2(ok328); unc-43(n498n1186)* double mutant exhibits a strong posterior neurite phenotype (49%, n=122), to a similar degree as *lin-44(n1792)* mutant (**Figures 3A, 3B, and 3D**).

**Figure 3.**
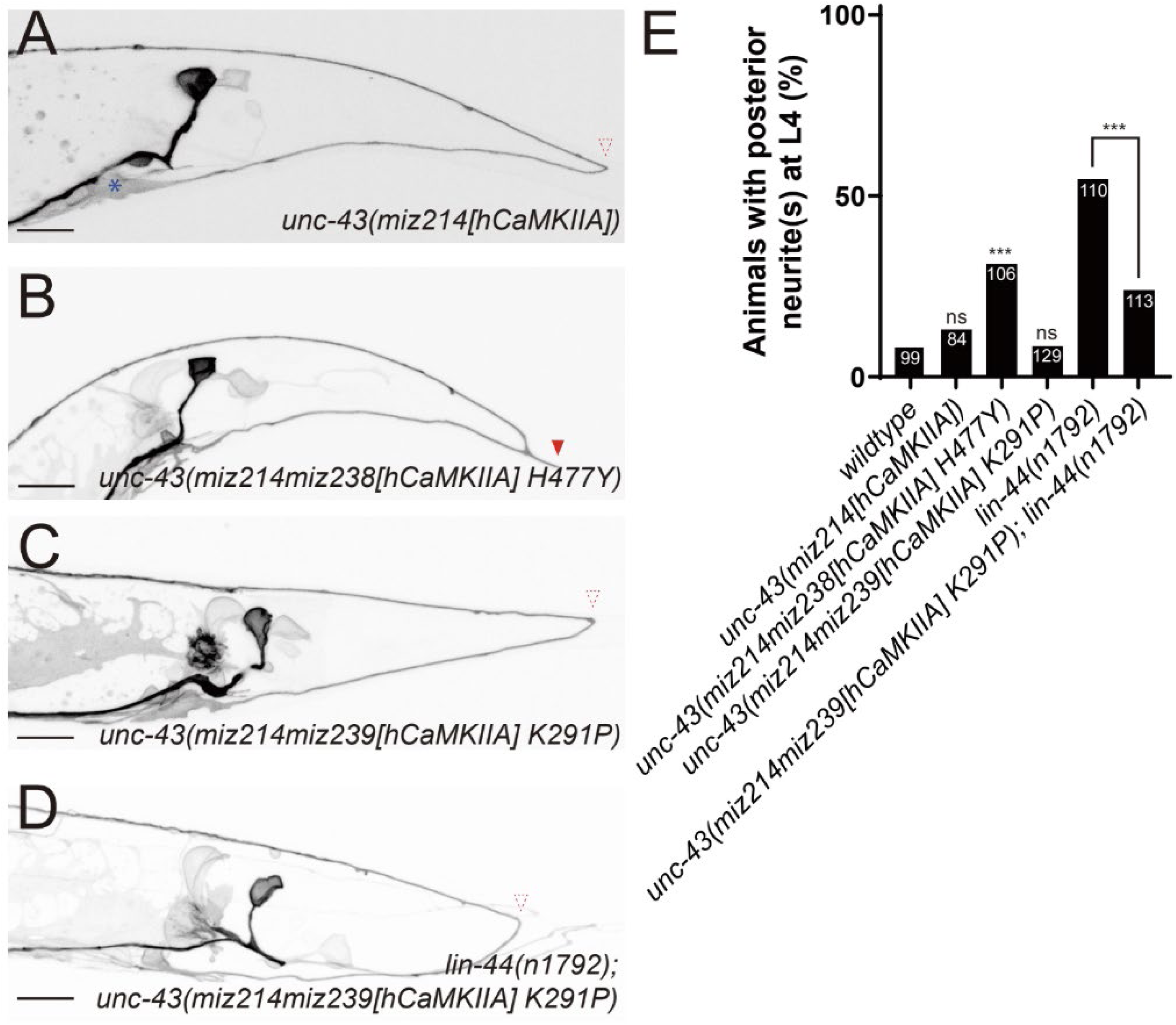
*pkc-2* functions in parallel to *unc-43* to regulate PDB neurite pruning. **(A-C)** Representative images of PDB of *pkc-2(ok328)* **(A)**, *unc-43(n498n1186); pkc-2(ok328)* double mutant **(B),** and *lin-44(n1792); unc-43(n498n1186); pkc-2(ok328)* triple mutant (**C**) at the L4 stage. Red arrowheads represent ectopic posterior neurites. Asterisks denote the PDB cell bodies. **(D)** Quantification of the animals with ectopic posterior neurites. Sample numbers are shown in each bar. ***p < 0.001; **p < 0.002; *p < 0.033; ns, not significant (Fisher’s exact test). Note the quantification of wildtype, *lin-44(n21792)* and *unc-43(n498n1186)* are from Figure 1G. Scale bars: 10 μm.

To confirm that the posterior neurites observed in *pkc-2(ok328)* mutants at the L4 stage are due to pruning defects, we examined the posterior neurites of PDB at the L2 and the L4 stages. We found that 30% (n = 57) of posterior neurites observed at the L2 stage failed to prune in *pkc-2(ok328)* mutants (**Figures 2C, 2E, and 2F**). In the *unc-43(n498n1186); pkc-2(ok328)* double mutants, 51% (n = 75) of posterior neurites present at the L2 stage fail to prune, a penetrance comparable to that of *lin-44(n1792)* mutant (**Figures 2D–2F**). Furthermore, *lin-44(n1792); unc-43(n498n1186); pkc-2(ok328)* triple mutants exhibit a similar degree of posterior neurite phenotype to that seen in *lin-44(n1792)* single mutant and *unc-43(n498n1186); pkc-2(ok328)* double mutant (**Figures 3C, 3D**). Taken together, these results suggest that *pkc-2* and *unc-43* function in parallel and act in the same pathway as *lin-44* in regulating neurite pruning.

We previously showed that, in addition to its role in neurite pruning, LIN-44/Wnt is also required for the PDB cell fate specification and neurite guidance (**Figure S3A**) (Lu & Mizumoto, 2019). In contrast, we did not observe significant defects in PDB cell fate specification and neurite guidance in the loss-of-function mutants of *unc-43* and *pkc-2.* Furthermore, the gain-of-function mutations of *unc-43* did not suppress the PDB cell fate specification and neurite guidance defects observed in *lin-44(n1792)* mutants (**Figures S3A, S3B**). These results indicate that Wnt-calcium signaling specifically regulates neurite pruning in PDB development.

### IP3 signaling and *itr-1/IP3R* are required for PDB neurite pruning

Both PKC-2/PKC and UNC-43/CaMKII are calcium-dependent kinases. The Wnt-calcium pathway is known to upregulate intracellular calcium through inositol 1,4,5-trisphosphate receptor (IP3R), ryanodine receptor (RyR), and Voltage-gated calcium channels (VGCCs) (LeBoeuf et al., 2020; Li et al., 2009; McQuate et al., 2017). We therefore examined the PDB neurite structure in the mutants of *itr-1/*IP3R (Baylis et al., 1999), *unc-68/*RyR (Maryon et al., 1996), *egl-19*/CaV1α, *unc-2*/CaV2α1, *unc-36*/CaV1α2, and *cca-1*/CaV3α (Lee et al., 1997; Schafer & Kenyon, 1995; Schafer et al., 1996; Steger et al., 2005). Among them, the temperature-sensitive mutant of *itr-1(sa73)* showed a significantly higher penetrance of posterior neurites at the L4 stage (33%, n = 82) compared to the wild-type animals (15%, n = 87) at the restrictive temperature 25 °C (**Figures 4A, 4B**). While it does not exclude the possibility that other calcium channels also contribute to neurite pruning, this result suggests that IP3R is the primary calcium channel in PDB neurite pruning. Similar to *unc-43* and *pkc-2* mutants, *itr-1(sa73)* mutants did not exhibit significant defects in PDB cell fate specification or neurite guidance at the restrictive temperature, indicating its specific role in neurite pruning (**Figure S3B**).

**Figure 4.**
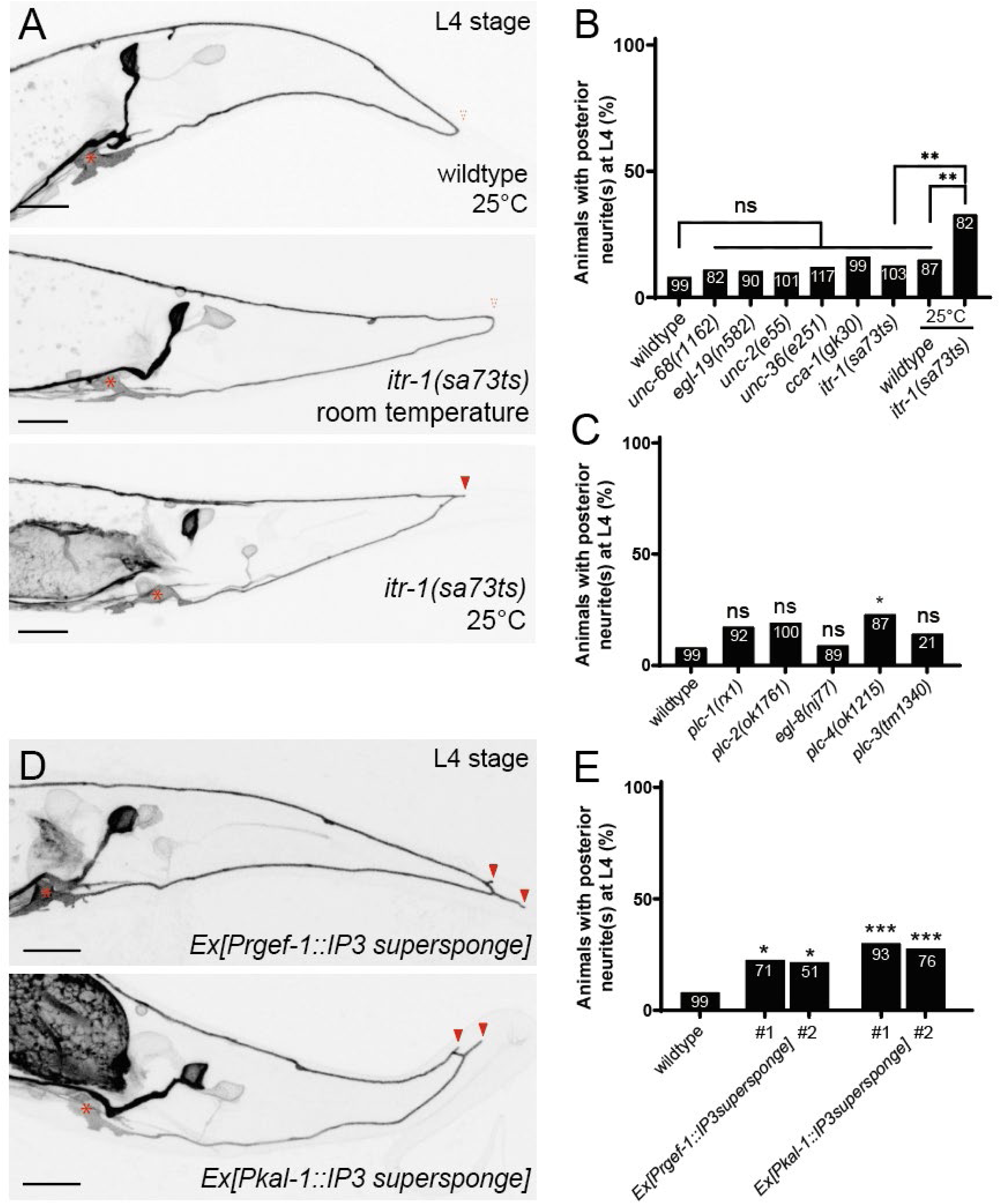
*itr-1* is required for PDB neurite pruning. **(A)** Representative images of PDB in wildtype (top), *itr-1(sa73ts)* mutant at the room temperature (middle), and *itr-1(sa73ts)* mutant at the restrictive temperature (bottom) at the L4 stage. Red arrowhead represents the ectopic posterior neurite, and dotted arrowheads represent the absence of posterior neurites. Asterisks denote the PDB cell body. **(B, C) (D)** Representative images of PDB in wildtype with IP3 supersponge overexpression from the *rgef-1* (top) and *kal-1* (bottom) promoters at the L4 stage. Red arrowheads represent the ectopic posterior neurites. **(E)** Quantification of the animals with ectopic posterior neurites. Sample numbers are shown in each bar. ***p < 0.001; **p < 0.002; *p < 0.033; ns, not significant (Fisher’s exact test). Note the quantification of wildtype is from Figure 1G. Scale bars: 10 μm.

IP3R is a ligand-gated calcium channel primarily localized on the endoplasmic reticulum (ER) membrane, where it releases calcium in response to cytoplasmic IP3 (Bezprozvanny et al., 1991). Phospholipase C (PLC) produces IP3 from Phosphatidylinositol 4,5-bisphosphate (PIP2). *C. elegans* possesses five PLCs: *egl-8*/PLCβ, *plc-2/*PLCβ-like, *plc-3*/PLCγ, *plc-4*/PLCδ, and *plc-1/*PLCε (Baylis & Vazquez-Manrique, 2012). Among them, *plc-4(ok1215)* mutants exhibited a moderate posterior neurite phenotype at the L4 stage (23%, n = 87) (**Figure 4C**). Furthermore, overexpression of an IP3 super-sponge, which binds to IP3 thereby blocking IP3 signaling (Ghosh-Roy et al., 2010; Walker et al., 2002), under the pan-neuronal (P*rgef-1*) or the PDB-specific (P*kal-1*) promoters led to posterior neurite phenotype at the L4 stage, suggesting that IP3 signaling is required cell-autonomously in PDB for neurite pruning (**Figures 4D, 4E**). Overexpression of the IP3 super-sponge did not enhance the posterior neurite phenotype of *pkc-2(ok328)* and *unc-43(n498n1186)* mutants (**Figure S4**), suggesting IP3 signaling function in the same pathway as the two kinases. Taken together, these results are consistent with our hypothesis that the two calcium-dependent kinases UNC-43 and PKC-2 are regulated by IP3-dependent calcium release.

### *lin-44*/Wnt regulates PDB neurite pruning through *itr-1/*IP3R

To test whether LIN-44/Wnt regulates neurite pruning through ITR-1/IP3R, we examined the posterior neurite phenotype in the *lin-44(n1792); itr-1(miz414)* double mutants. We generated the gain-of-function allele of *itr-1(miz414)*, which carries the same mutation (R511C) as the known gain-of-function allele *itr-1(sy290)* (Clandinin et al., 1998). We found that *itr-1(miz414)* suppressed the posterior neurite phenotype (**Figures 5A, 5B**), but not the cell fate specification and neurite guidance defects of *lin-44(n1792)* mutant (**Figure S3B**). This result is consistent with our hypothesis that *itr-1* functions downstream of *lin-44* to regulate neurite pruning.

**Figure 5.**
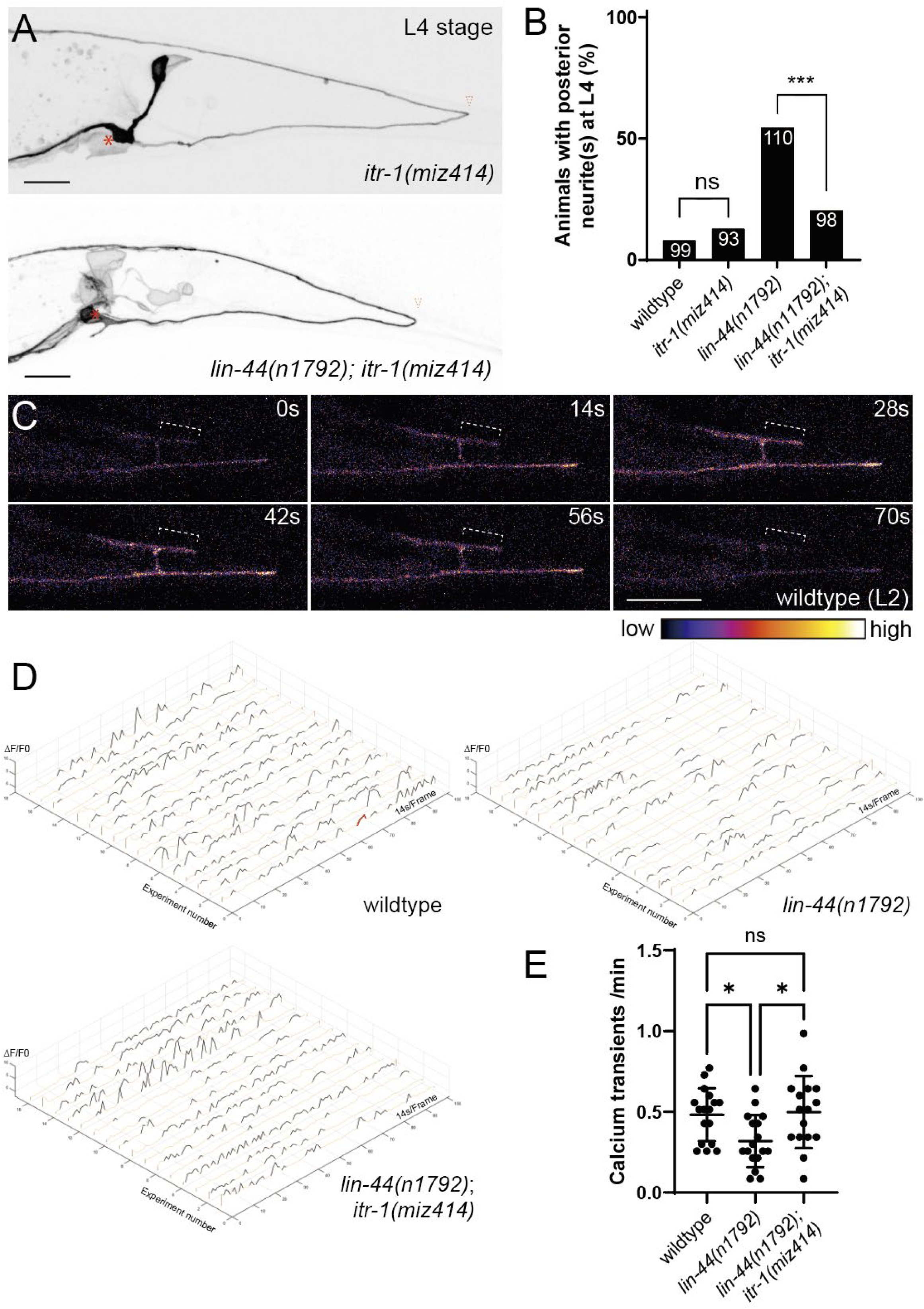
*itr-1* functions downstream of *lin-44* to regulate PDB neurite pruning. **(A)** Representative images of PDB in *it2r-1(miz414)* (top) and *lin-44(n1792); itr-1(miz414)* double mutant (bottom) at the L4 stage. Red arrowheads represent ectopic posterior neurites, and dotted arrowheads repr2esent the absence of posterior neurites. Asterisks denote the PDB cell body. **(B)** Quantification of animals with ectopic posterior neurites. Sample numbers are shown in each bar. ***p < 0.001; ns, not significant (Fisher’s exact test). Note that the quantifications of wildtype *and lin-44(n1792)* are from Figure 1G. Scale bars: 10 μm. **(C)** Heatmap of a time-lapse image showing one calcium transient event of a wildtype animal at the L2 stage. Dotted lines denote the segment of neurite that is set as the region of interest (ROI). **(D)** Change in GCaMP7s fluorescence within the ROI represented by ΔF/F0 in wildtype (top left), *lin-44(n1792)* (top right), and *lin-44(n1792)*; *itr-1(miz414)* (bottom) animals at the L2 stage. Black segments of the plot denote the recognized calcium transients, and the red dots denote the calcium transient shown in panel C. **(E)** Plot of calcium transient frequencies in wildtype, *lin-44(n1792)*, and *lin-44(n1792)*; *itr-1(miz414)* mutants. *p < 0.033; ns, not significant (one-way ANOVA test).

We next examined calcium dynamics during PDB neurite pruning using the genetically encoded calcium indicator, GCaMP7s (Dana et al., 2019). In wild-type animals, we observed calcium transients along the PDB neurite, including the posterior neurites, at the L2 stage when neurite pruning occurs (**Figure 5C**). We then asked whether the calcium transient is *lin-44*/Wnt dependent. We quantified the frequency of calcium peaks that are at least 100% higher than the baseline (see Methods) in 17 animals of wildtype and *lin-44(n1792)* and found that *lin-44(n1792)* mutant animals exhibit significantly lower calcium transient frequency (0.32/minute) compared to wildtype animals (0.48/minute) (**Figures 5D, 5E**). Consistent with the hypothesis that *itr-1* functions downstream of *lin-44* to regulate calcium signaling, *itr-1(miz414)* suppressed the phenotype of reduced calcium transients in *lin-44(n1792)* mutants and exhibited a normal calcium transient frequency (0.50/minute) (**Figure 5D, 5E**). It is worth noting that these calcium transients are specific during the L2-L3 stage when neurite pruning occurs. At the L4 stage, when pruning has completed, the calcium transients are nearly absent in wildtype, *lin-44(n1792),* and *itr-1(miz414)* mutant animals (**Figure S5**).

As ITR-1/IP3R is an ER-localized calcium channel, we examined the subcellular localization of ER using the GFPnovo2-tagged ER-resident signal peptidase SP12 (Liu et al., 2019). We found that the ER is extended into the posterior neurites during the L2-L3 stage in both wild-type and *lin-44(n1792)* mutant (**Figure S6**).

### *pkc-2*/PKC regulates clathrin-dependent endocytosis to regulate PDB neurite pruning

Clathrin-dependent endocytosis has been shown to play a role in removing membrane components from the growth cone in response to a calcium signal (Tojima et al., 2010, 2014). As the pruning of PDB neurites is mediated by retraction (Lu & Mizumoto, 2019), we examined the involvement of clathrin-dependent endocytosis in PDB neurite pruning. Among the core components of the clathrin-dependent endocytic pathway we examined (*unc-101/*AP-1µ, *apa-2*/AP-2α, *apb-3*/AP-3β, *apm-3*/AP-3µ and *chc-1/*clathrin heavy chain) (Koushika & Nonet, 2000; Lee et al., 1994; Shim & Lee, 2005), the null mutants *unc-101(m1)* and *apa-2(miz461)* exhibit significant posterior neurite phenotype at the L4 stage (**Figures 6A, 6B**). The temperature-sensitive mutant of clathrin heavy chain, *chc-1(b1025)*, also exhibits significant posterior neurite phenotypes at the restrictive temperature at the L4 stage (**Figures 6A, 6B**).

**Figure 6.**
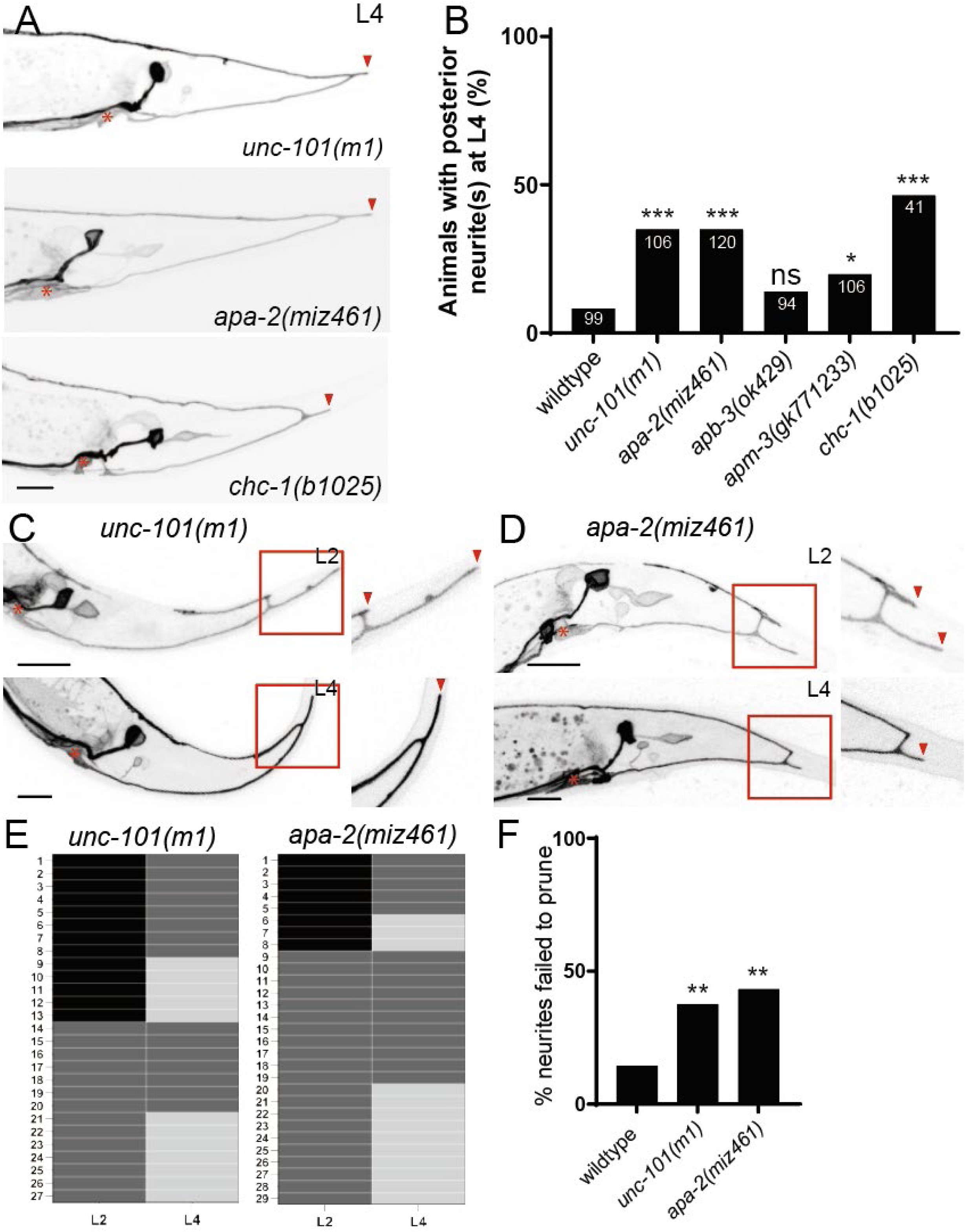
Clathrin-dependent endocytosis is required for PDB neurite pruning. **(A)** Representative images of PDB at the L4 stage of *unc-101(m1)* (top), *apa-2(miz461)* (middle), and *chc-1(b1025)* (bottom). Red arrowheads represent ectopic posterior neurites. Asterisks denote the PDB cell bodies. **(B)** Quantification of animals with ectopic posterior neurites. Sample numbers are shown in each bar. ***p < 0.001; *p < 0.033; ns, not significant (Fisher’s exact test). Note the quantification of wildtype is from Figure 1G. **(C, D)** Representative images of single animals of *unc-101(m1)* (**C**) and *apa-2(miz461)* (**D**) at L2 (top panels) and L4 stages (bottom panels). Red arrowheads indicate the posterior neurites. Asterisks denote the PDB cell body. **(E)** Quantification of the posterior neurite number of individual animals at L2 and L4 stages in each genetic background. **(F)** Quantification of the posterior neurite pruning frequency. **p < 0.002 (Fisher’s exact test). All scale bars: 10 μm.

To determine whether these mutants have neurite pruning defects, we examined pruning frequency by assessing PDB morphology at the L2 and L4 stages. In *unc-101(m1)* and *apa-2(miz461)* mutants, 38% (n=40) and 43% (n=37) of posterior neurites present at the L2 stage failed to prune, respectively (**Figures 6C, 6D**). These results indicate that clathrin-dependent endocytosis is required for neurite pruning in PDB.

Consistent with the involvement of endocytic pathway components in neurite pruning, we often observed puncta of an early endosome marker, RAB-5 (Sato et al., 2005), in the posterior neurite of wild-type animals (79%, n = 24) at the L2 stage when neurite pruning occurs (**Figure 7A**). The proportion of animals with RAB-5 puncta at the posterior neurite was significantly lower in *unc-101* (40 %, n = 20) and *apa-2* (31 %, n = 22) mutants (**Figures 7B, 7C, and 7G**). Interestingly, we often observed retrograde transport of RAB-5 puncta along the posterior neurite (**Figure 7H**), which may reflect membrane-removal events during neurite pruning.

**Figure 7.**
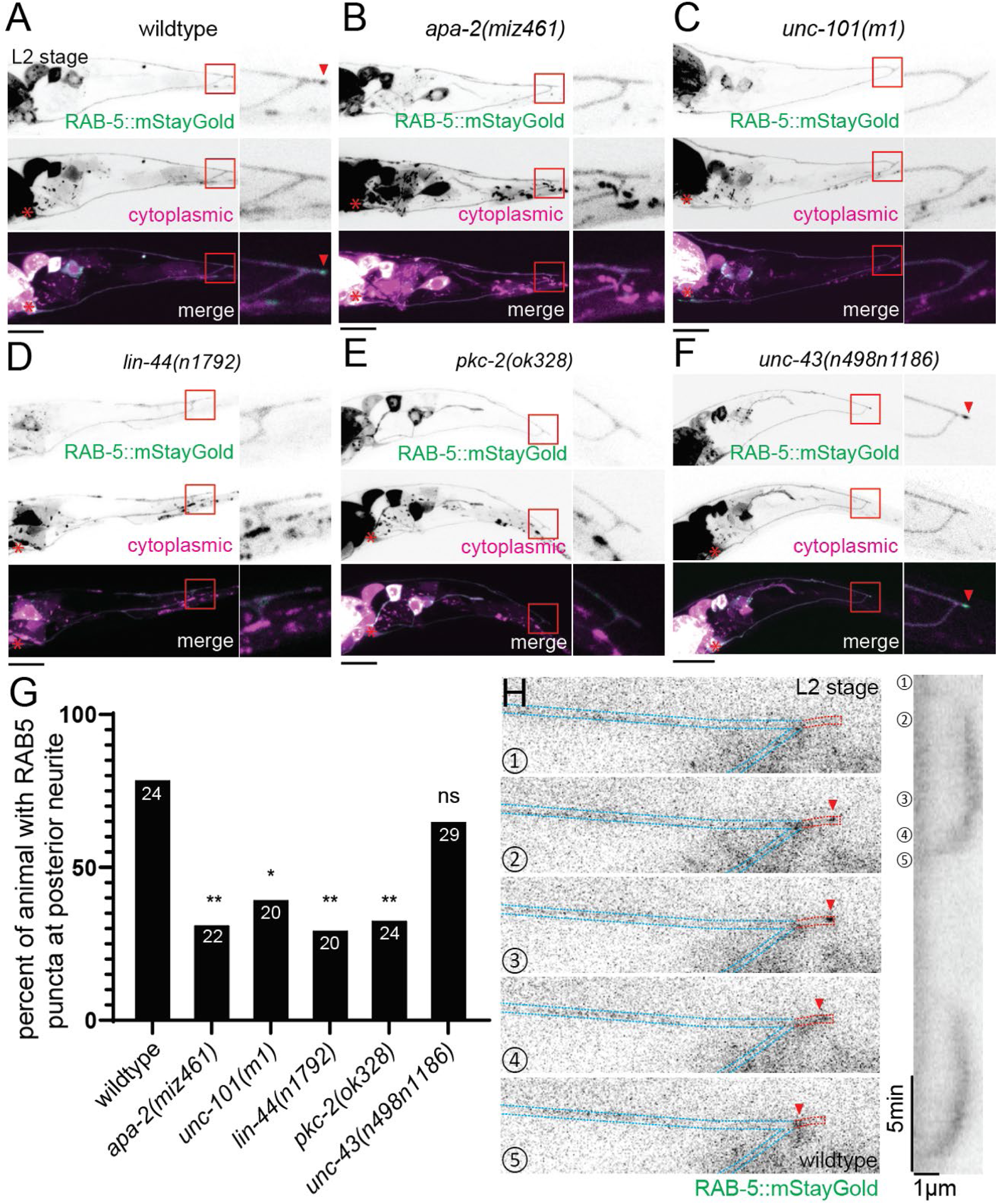
RAB-5 localization at the posterior neurites at L2 stage depends on *lin-44* and *pkc-2*. **(A-F)** Representative images of RAB-5::mStayGold localization (top panels), PDB neurite structure labeled with cytoplasmic mScarlet (middle panels) and merged images (bottom panels) of wild type (**A**), *apa-2(miz461)* (**B**), *unc-101(m1)* (**C**), *lin-44(n1792)* (**D**), *pkc-2(ok328)* (**E**) and *unc-43(n498n1186)* (**F**). Red arrowheads indicate the RAB-5 puncta at the posterior neurites. Red boxes indicate the region of magnified images. Scale bars: 10 μm. **(G)** Quantification of the percentage of animals with RAB-5 puncta at the posterior neurite of PDB. Sample numbers are shown in each bar. **p < 0.002; *p < 0.033; ns, not significant (Fisher’s exact test). **(H)** Time-lapse and kymograph showing the retrograde movement of RAB-5 punctum. The red dotted line represents the ROI for the kymograph, and the red arrowheads indicate the RAB-5 punctum forming at the tip of the posterior neurite and undergoing retrograde movement (bottom). Scale bar: 1 μm.

Clathrin-dependent Frizzled endocytosis is crucial for proper activation/inactivation of Wnt signaling (Brunt & Scholpp, 2018). Given that LIN-17/Frizzled is enriched in the pruning neurites (**Figure 1H**) (Lu & Mizumoto, 2019), the neurite pruning defects observed in the endocytic pathway mutants may be attributed to disrupted LIN-17 endocytosis. However, the LIN-17::GFP puncta in the posterior neurites are unaffected in the *apa-2* mutant (**Figure 8A**), suggesting that the endocytic pathway does not regulate neurite pruning via LIN-17 endocytosis.

**Figure 8.**
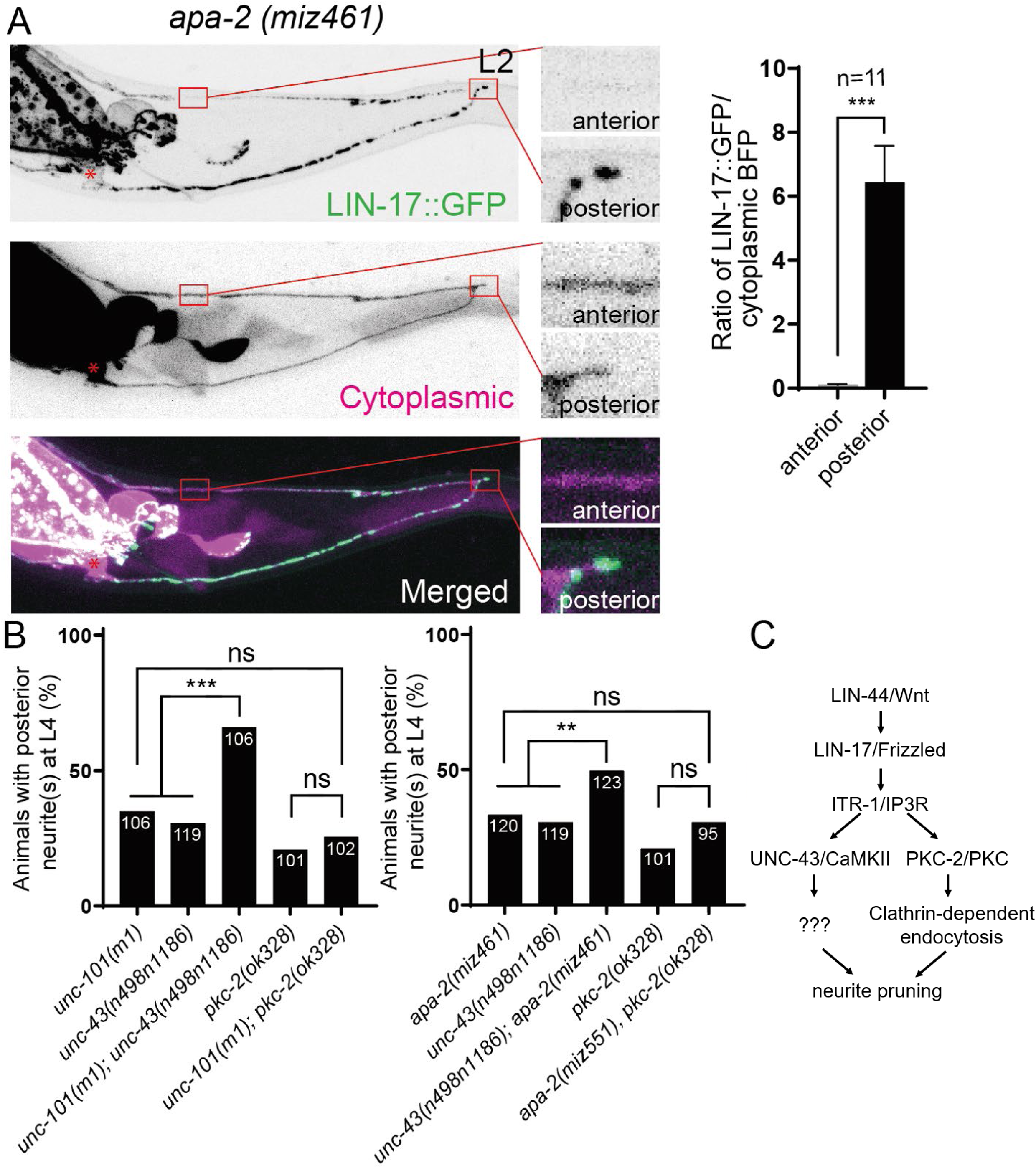
*pkc-2* but not *unc-43* regulates clathrin-dependent endocytosis during neurite pruning. **(A)** Representative images of LIN-17::GFP localization (top panels), PDB neurite structure labeled with cytoplasmic BFP (middle panels) and merged images (bottom panels) in *apa-2(miz414)* animals. Magnified images of anterior and posterior neurites are also shown. Asterisks represent the PDB cell body. Quantification of the normalized GFP/BFP ratio at the anterior and posterior growth cones for each genotype is shown on the right. Error bars indicate mean ± SEM. ***p < 0.001 (Ratio paired t-test). **(B)** Qu2antification of animals with ectopic posterior neurites. Sample numbers are shown in each bar. Note the quantifications of *unc-43(n498n1186)* and *pkc-2(ok328)* are from Figure 1 and Figure 3. ***p < 0.001; **p < 0.002; ns, not significant (Fisher’s exact test). Scale bars: 10 μm. **(C)** The proposed model of Wnt-calcium pathway regulating neurite pruning in PDB posterior neurites.

Lastly, we asked whether Wnt signaling regulates clathrin-dependent endocytosis. The RAB-5 puncta in the posterior neurites are greatly diminished in *lin-44(n1792)* and *pkc-2(ok328)* mutants, while they are unaffected in *unc-43(n498n1186)* mutants (Figure 7D-7G), suggesting that endocytosis is regulated by the Wnt-PKC-2 axis but not by UNC-43. Consistently, the *apa-1(miz461)* and *unc-101(m1)* enhance the posterior neurite phenotype of *unc-43(n498n1186)* but not *pkc-2(ok328)* mutants (**Figure 8B**). This result indicates that *apa-2* and *unc-101* function in the same genetic pathway as *pkc-2* and in parallel with *unc-43*. Taken together, our data show that LIN-44/Wnt regulates neurite pruning via PKC-2 and UNC-43, and the clathrin-dependent endocytic pathway acts downstream of PKC-2 (**Figure 8C**).

## Discussion

Developmental neurite pruning is a critical process in shaping functional neural circuits, and its dysregulation has been implicated in various neurodevelopmental disorders. Specifically, conditions such as autism spectrum disorders (ASD) and schizophrenia are associated with aberrant and deficient neuronal connections during adolescence, respectively, which may result from misregulated neurite and/or synapse pruning (Penzes et al., 2011; Riccomagno & Kolodkin, 2015). In this study, we show that Wnt-calcium signaling regulates neurite pruning in *C. elegans*. Specifically, Wnt acts through the calcium channel, IP3R, and two calcium-dependent kinases, CaMKII and PKC. Furthermore, we identify clathrin-mediated endocytosis as a downstream effector of PKC. Importantly, missense variants in the PKC and CaMKII have been identified in patients with neurodevelopmental disorders, including ASD and intellectual disabilities (Chia et al., 2018; Coe et al., 2019; De Rubeis et al., 2014; Kury et al., 2017; Pan et al., 2025; Proietti Onori et al., 2018; Rigter et al., 2024). Moreover, a growing piece of evidence suggests the involvement of endocytic pathways in neurodevelopmental disorders (Baschieri et al., 2020; Helbig et al., 2019; Stankovic et al., 2025). We showed that humanized *C. elegans* carrying human CaMKIIA can fully replace the function of UNC-43 in neurite pruning, and mutations found in patients with intellectual disability cause neurite pruning defects. Combined with the latest genome-editing technologies and the highly conserved genetics between *C. elegans* and humans in neurodevelopment, our present work affirms the utility of *C. elegans* genetics in the study of fundamental processes of normal neurodevelopment and disease conditions.

In addition to neurite pruning, Wnt signaling regulates other aspects of PDB development, including cell fate specification and axon guidance (Jiang & Sternberg, 1998; Lu & Mizumoto, 2019). We also showed that, by using membrane-tethered LIN-44, the gradient distribution of LIN-44 is dispensable for PDB neurite pruning but necessary for PDB synapse patterning, cell fate specification, and axon guidance (Lu & Mizumoto, 2019). In this study, we found that none of the mutants of Wnt-calcium signaling components exhibited defects in cell fate specification or axon guidance in PDB. Therefore, it is likely that gradient-independent Wnt signaling specifically activates the Wnt-calcium pathway to induce neurite pruning. One possible mechanism is that different downstream cascades require distinct Wnt concentration thresholds, and that Wnt-calcium signaling requires the highest Wnt concentration, which can only be achieved in close proximity to LIN-44-expressing cells. Early studies of neurite pruning showed that this process often occurs within a defined developmental time window (Bagri et al., 2003; Walsh & Lichtman, 2003). Similarly, PDB neurite pruning starts at the L2 stage and is completed by the L4 stage. Consistently, we found that calcium transients are present during the L2–L3 stages when pruning occurs, but are nearly absent at the L4 stage. The temporal correlation between calcium transients and neurite pruning suggests that pruning is tightly regulated within a specific developmental time window. PDB neurite pruning occurs prior to synapse formation and is independent of neurotransmitter release (Lu & Mizumoto, 2019). It is possible that calcium transients are increased only when posterior neurites grow towards Wnt-expressing cells at the L2 stage to exceed the Wnt threshold to activate the Wnt-calcium pathway. However, the mechanisms behind the downregulation of calcium signaling after the L4 stage remain unknown.

We observed only a mild reduction of calcium transients in *lin-44*/Wnt mutant animals. This is consistent with the incomplete penetrance of PDB neurite pruning defects (48%) in *lin-44* mutants. It is possible that Wnt increases the frequency of calcium transients via IP3R, thereby increasing the probability of activating CaMKII and PKC to induce neurite pruning. This also implies that additional signaling pathways or calcium channels may contribute to calcium dynamics in PDB. Screening of additional mutants with PDB neurite pruning defects will reveal the complexity of developmental neurite pruning mechanisms.

Clathrin-mediated endocytosis is proposed to retrieve membrane components during growth cone repulsion in the Drosophila C4da neuron and chick dorsal root ganglion (Peng et al., 2015; Tojima et al., 2010). This has led to the hypothesis that endocytosis may also contribute to neurite pruning (Bodakuntla et al., 2021; Tojima & Kamiguchi, 2015). The involvement of clathrin-mediated endocytosis in PDB neurite pruning is consistent with a model in which it functions to remove excessive membrane components from retracting neurites. Our genetic data indicate that the clathrin-dependent pathway acts downstream of *pkc-2* but not of *unc-43*. Consistently, mammalian PLC and PKC regulate clathrin-dependent endocytosis via synaptojanin1 (Delos Santos et al., 2017). It is therefore interesting to examine the functions of *unc-26*, the sole ortholog of synaptojanin1 in *C. elegans*, in PDB neurite pruning.

In this study, we did not find the mechanisms downstream of *unc-43/CaMKII.* Mammalian CaMKII proteins are known to regulate actin and microtubule cytoskeletal scaffolds and their regulators (McVicker et al., 2015), including Cdk5 (Cyclin-dependent kinase 5) and Dbn1 (Drebrin 1) (Dhavan et al., 2002; Yamazaki et al., 2024). However, the embryonic lethality associated with loss of core cytoskeletal components and regulators prevented us from examining their roles in PDB neurite pruning. Future studies using the auxin-inducible degron system to overcome embryonic lethal phenotypes will be essential for determining how cytoskeletal regulation downstream of CaMKII contributes to neurite pruning.

## Materials and methods

### C. elegans genetics

The Bristol N2 strain was used as a wild-type reference. Animals were cultured in the nematode growth medium (NGM) with *Escherichia coli* strain OP50 as previously described standard protocols (Brenner, 1974), and maintained at room temperature (22°C) unless otherwise specified. All animals analyzed were hermaphrodites. The following allele were used in this study: *lin-44(n1792)* I, *apb-3(ok429)* I, *unc-101(m1)* I, *plc-3(tm1340)/mIn1 [mIs14 dpy-10(e128)]* II, *unc-36(e251)* III, *chc-1(b1025)* III, *egl-19(n582)* IV, *plc-4(ok1215)* IV, *itr-1(sa73)* IV, *itr-1(miz414)* IV, *unc-43(n498n1186)* IV, *unc-43(n498)* IV, *unc-43(miz521 [7xGFP11::unc-43])* IV, *unc-43(miz214[hCaMKIIA])* IV, *unc-43(miz214miz238[hCaMKIIA H477Y])* IV, *unc-43(miz214miz209[hCaMKIIA K291P])* IV, *egl-8(nj77)* V, *unc-68(r1162)* V, *plc-2(ok1761)* V, *apm-3(gk771233)* X, *unc-2(e55)* X, *cca-1(gk30)* X, *apa-2(miz461)* X, *apa-2(miz551)* X, *pkc-2(ok328)* X, *plc-1(rx1)* X. Genotyping primers are listed in the Key Resources Table.

### Plasmid construction

*C. elegans* expression plasmids were constructed using a derivative of pPD49.26 (A. Fire), the pSM vector. cDNAs of *unc-43*, IP3 sponge, SP12, and *rab-5* were obtained by RT-PCR from N2 mRNA using Superscript III First-strand synthesis system and Phusion High-Fidelity DNA Polymerase (Thermo Fisher Scientific). The codon-optimized GCaMP7s was synthesized using the GeneArt Synthesis service (Dana et al., 2019; Zhang et al., 2024) (Thermo Fisher Scientific). These cDNA constructs were then cloned into the *Asc*I and *Kpn*I sites of plasmid vectors containing P*kal-1* or P*rgef-1* (Thermo Fisher Scientific). The IP3 super-sponge was generated by a site-directed mutagenesis to introduce the same mutation as the R511C mutation in *itr-1(sy290)* (the sequences of IP3 sponge and super-sponge are kind gifts from Dr. A. Chisholm). The plasmid sequences were confirmed by Sanger sequencing (GENEWIZ from Azenta Life Sciences). The primer sequences used to construct these plasmids are available in the key resource table.

### Transgenes

The transgenic extrachromosomal arrays used in this study were generated using the standard microinjection method (Fire, 1986; Mello et al., 1991).

Tissue-specific rescue of *unc-43(n498n1186)* mutant phenotype: *mizEx610/mizEx611 [Prgef-1::unc-43* 10ng/µL*, Podr-1::RFP* 30ng/µL*], mizEx701/mizEx702 [Pkal-1::unc-43* 10ng/µL*, Podr-1::RFP* 30ng/µL]; IP3 super-sponge expression: *mizEx637/mizEx638 [Pkal-1::IP3 super-sponge* 10ng/µL*, Podr-1::RFP* 30ng/µL*], mizEx642/mizEx643 [Prgef-1::IP3 super-sponge* 10ng/µL*, Podr-1::RFP* 30ng/µL]; localization of LIN-17 in PDB: *mizEx295 [Pkal-1::lin-17::GFPnovo2* 5ng/µL, *Pkal-1::BFP* 30ng/µL, *Podr-1::GFP* 20ng/µL] (Lu & Mizumoto, 2019); localization of ER in PDB: *mizEx592[Pkal-1::GFPnovo2::SP12* 10ng/µL*, Pkal-1::mScarlet* 15ng/µL*, Podr-1::GFP* 20ng/µL]; localization of early endosomes in PDB: *mizEx633[Pkal-1::mStayGold::rab-5* 10ng/µL*, Pkal-1::mScarlet* 15ng/µL*, Podr-1::GFP* 20ng/µL].

The integrated transgene *mizIs39[Pkal-1::GCaMP7s* 20ng/µL*, Pkal-1::mScarlet* 20ng/µL] was generated by UV mediated transgene integration (Mariol et al., 2013). *mizIs64* [*Pkal-1::GFP1-10* 10ng/µL; *Podr-1::GFP* 20ng/µL] was generated by using the Fluorescent Landmark Interference (FLInt) method (Malaiwong et al., 2023). *mizSi27 [Peft-3::TdTomato]* at *oxTi365 V* was used as the FLInt destination (Lu and Mizumoto, submitted).

### CRISPR/Cas9 genome editing

Genome editing was performed using the CRISPR/Cas9 system according to the protocol described previously (Ghanta & Mello, 2020; Kurashina & Mizumoto, 2023). The alleles *itr-1(miz414), apa-2 (miz461)*, *apa-2(miz551), unc-43(miz521[7xGFP11::unc-43]), unc-43(miz214[hCaMKIIA]), unc-43(miz214miz209[hCaMKIIA K291P])* and *unc-43(miz214miz238[hCaMKIIA H477Y])* were generated using homology-dependent repair (HDR). Donor templates for *unc-43(miz521 [7xGFP11::unc-43])* was generated by amplifying tandem repeats of *GFP_11_* (He et al., 2019; Kurashina et al., 2025) flanked by 50-60 bp of homology arm sequences to the *unc-43* locus using Phusion DNA polymerase (New England Biolabs). The codon-optimized human CaMKIIA sequence with synthetic introns was synthesized using GeneArt Gene Synthesis (Thermo Fisher Scientific) and cloned into the plasmid containing 500 bp of homology upstream and downstream of the *unc-43* locus, flanking a dual selection cassette (Au et al., 2019) to generate the *unc-43(miz214[hCaMKIIA])* donor template plasmid. Sequences of ssODNs for the *itr-1* and *apa-2* alleles are included in the supplemental materials. The donor templates were co-injected with a co-injection marker plasmid and Cas9 ribonucleoprotein complex (Integrated DNA Technologies). F1s with the co-injection marker were singled and screened for heterozygote mutation by PCR genotyping, and homozygote F2s without the co-injection marker were later selected. The edits were confirmed by Sanger sequencing (GENEWIZ from Azenta Life Sciences). The sequence of gRNA, primer sequences for amplifying the HDR templates, DNA oligos used for HDR templates, and primer sequences for genotyping are in the supplemental materials.

### Confocal microscopy

Images of fluorescently-labeled neurons and proteins were captured in live *C. elegans* using a Zeiss LSM800 Airyscan confocal microscope (Carl Zeiss, Germany). Animals at the L2 and L3 stage are imaged with a 63x/1.4 oil DIC objective lens; all other stages are imaged with the 40x/1.3 oil objective lens. Worms were immobilized on 2% agarose pad using a mixture of 7.5 mM levamisole (Sigma-Aldrich) and 0.225M 2,3-butanedione monoxime (BDM) (Sigma-Aldrich). For time-lapse imaging, ∼20 animals were mounted onto 10% agarose pads prepared with M9 buffer and immobilized using 0.5 µL of 2.5 wt% 0.10 µm polystyrene latex microspheres (Alfa Aesar # 427124Y). The coverslip was sealed with Vaseline (Vaseline Jelly Original) to prevent dehydration of the animals and agarose pad during imaging. Images were processed by z-stack maximum projection using Zen software (Carl Zeiss) and analyzed with ImageJ (NIH, USA). To quantify LIN-17::GFP signal at the posterior neurite and anterior growth cone, a 0.5 µm^2^ region of interest (ROI) was defined at each tip of the two neurites. For background correction, the average pixel intensity of the GFP and BFP channels was measured from adjacent non-neuronal regions using ROIs of identical size. These background values were subtracted from the corresponding GFP and BFP intensities measured at the neurite tips. The corrected GFP signal was then normalized by calculating the ratio of the average GFP to BFP pixel intensities within each ROI.

For GCaMP7s imaging, we acquired 5 z-stacks every 14 seconds, and an ROI of a 0.5 µm-wide line along the dorsal posterior neurite at the L2 stage was used to quantify the mean signal intensity. The first quartile of measurement was defined as baseline F_0_. Normalized fluorescence was calculated using the formula ΔF/F_0_ (ΔF= F_current_ - F_0_). Peaks of calcium transients were then identified and quantified using PeakCaller (Artimovich et al., 2017).

### Semi-synchronization of *C. elegans*

Gravid adults were bleached using 2.5% sodium hypochlorite and 0.5□N sodium hydroxide for 5□min to isolate eggs. Eggs were washed twice with M9 buffer and seeded onto the NGM plates. To observe pruning, animals with posterior neurites were rescued from the slides after imaging at the L2 stage (∼26□hours post-seeding), and transferred to individual NGM plates to resume their development for 24□hours before re-imaging.

### Quantification of ectopic posterior neurites

The phenotype of posterior neurites in PDB at the L4 stage was quantified using Zeiss AxioPlan2 Fluorescence compound microscope with 40x/1.3 oil objective lens (Carl Zeiss, Germany). Around 30-40 animals at the L4 stage were immobilized on a 2% agarose pad using a mixture of 7.5 mM levamisole (Sigma-Aldrich) and 0.225M BDM (Sigma-Aldrich).

### Statistics

Data was analyzed using Prism10.4.0 (GraphPad Software, USA) and JupyterLab. Fisher’s exact test with the Holm–Bonferroni method was used to compare the penetrance of the ectopic posterior neurite phenotype at the L4 stage and the frequency of neurite pruning. Ratio paired t-test was used to compare normalized LIN-17::GFP between the anterior and posterior neurites. One-way ANOVA with Tukey’s multiple comparison test was used to compare the frequency of calcium transients. *, ** and *** represent p-value <0.05, <0.01 and <0.001 respectively. The sample size is indicated in the figures.

## Acknowledgements

We thank Dr. Don Moerman for his guidance and support throughout the work, and for sharing lab equipment and reagents, Dr. Kurt Haas, Dr. Ben Matthews, and Dr. Kenji Sugioka for critical feedback, and Dr. Callista Yee for comments on the manuscript. We thank Dr. A. Chisholm for kindly sharing the IP3 super-sponge plasmid sequence. We also thank former and current members of the Mizumoto Lab for general discussions and comments on the manuscript, Ilham Hakim for assistance in generating the *itr-1(miz414)* allele, and Dr. Ardalan Hendi for primer design for genotyping *apa-3* and *apm-3* mutants. Some strains used in this study were obtained from the Caenorhabditis Genetics Center (CGC), which is funded by the NIH Office of Research Infrastructure Programs (P40 OD010440), and the National Bioresource Project (Japan). This project is funded by the Natural Sciences and Engineering Research Council of Canada (RGPIN2015–04022, RGPIN-2021-03154) and the Canadian Institutes of Health Research (OGB-190360, PJT-191897). ML was supported by UBC Cell 1-year fellowship, Michael Smith Memorial Fellowship, David W. Strangway Fellowship and Josephine T. Berthier Fellowship. MK is supported by a UBC 4-year fellowship and CIHR CGRS-D. KM was supported by a Tier 2 Canada Research Chair program.

## Primer sequences

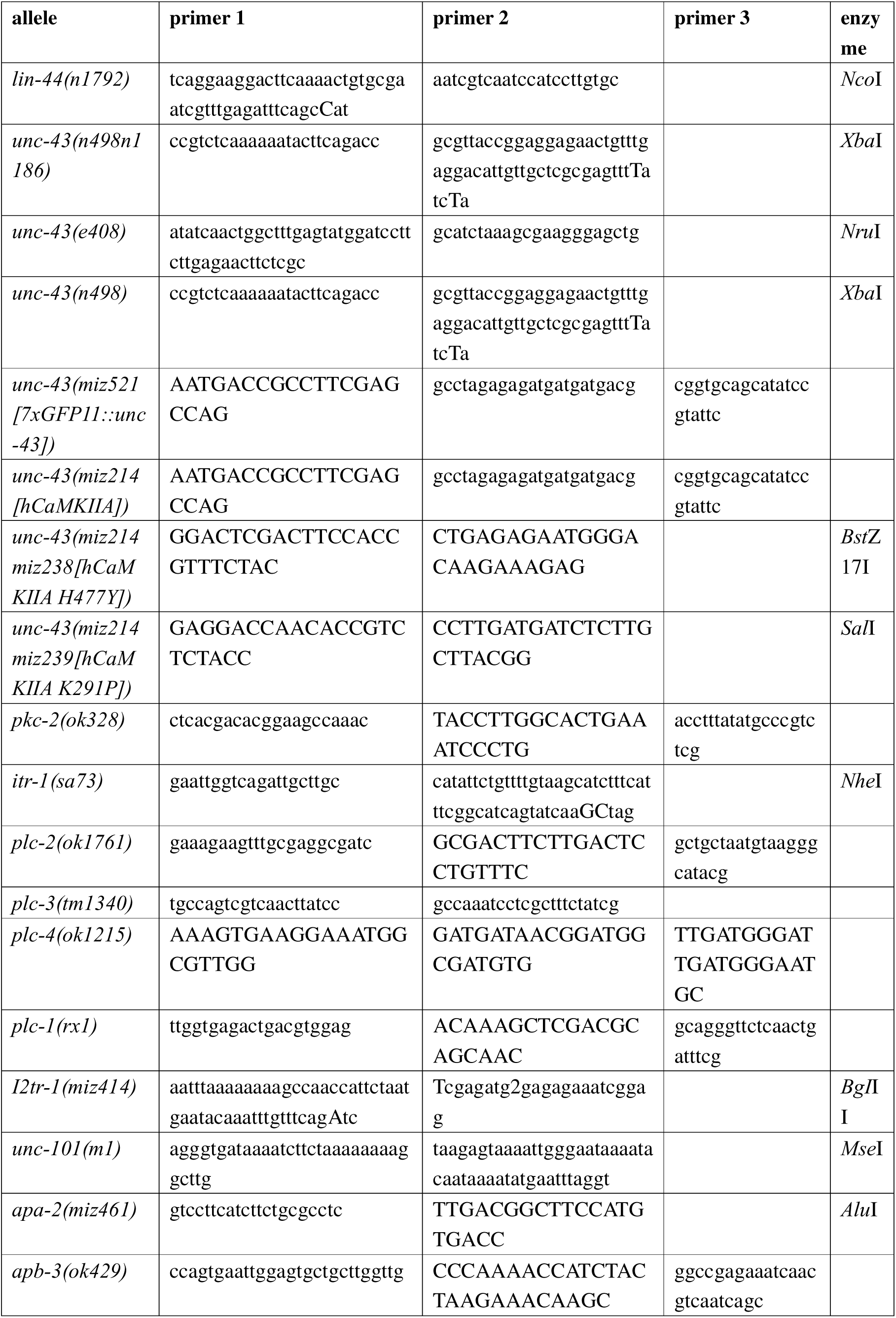
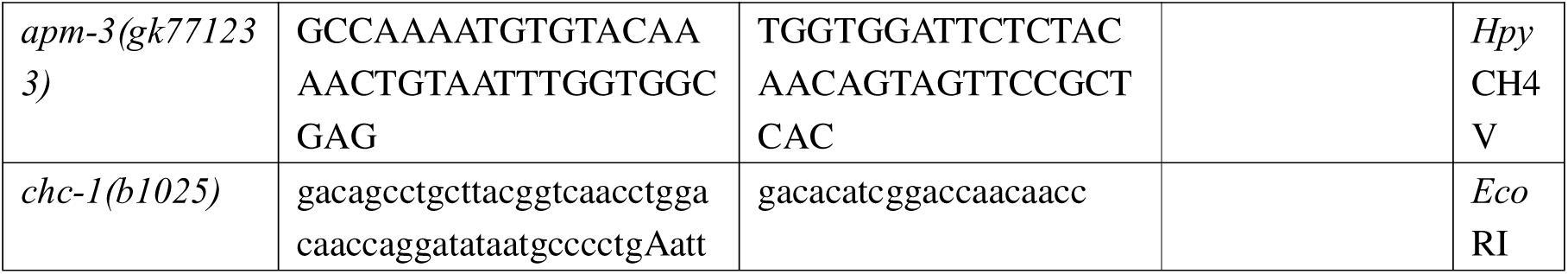

## CRISPR/Cas9 gRNA and repair HDR sequences

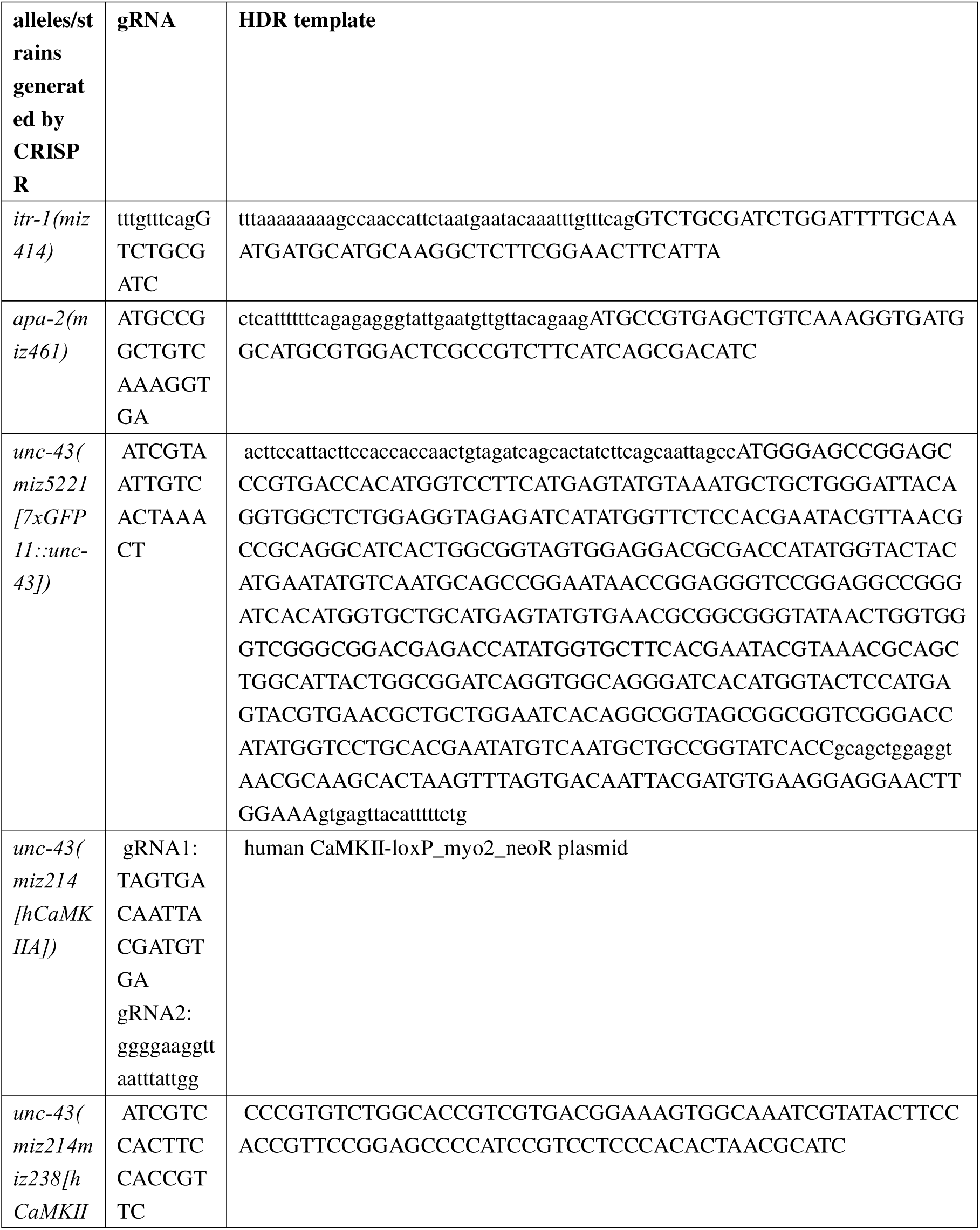
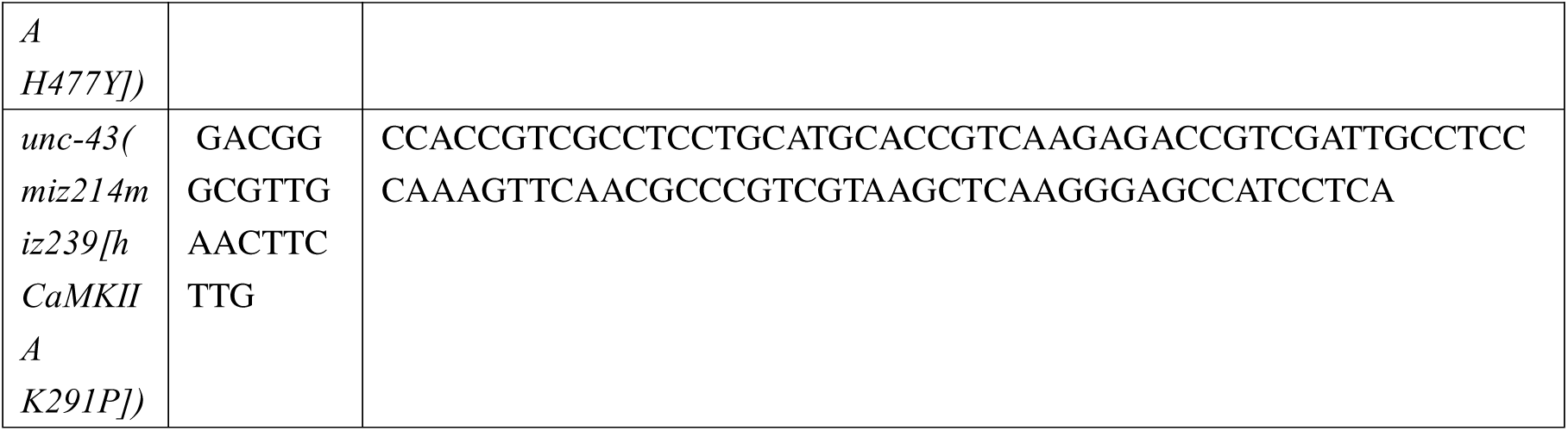

---

**Figure S1.**
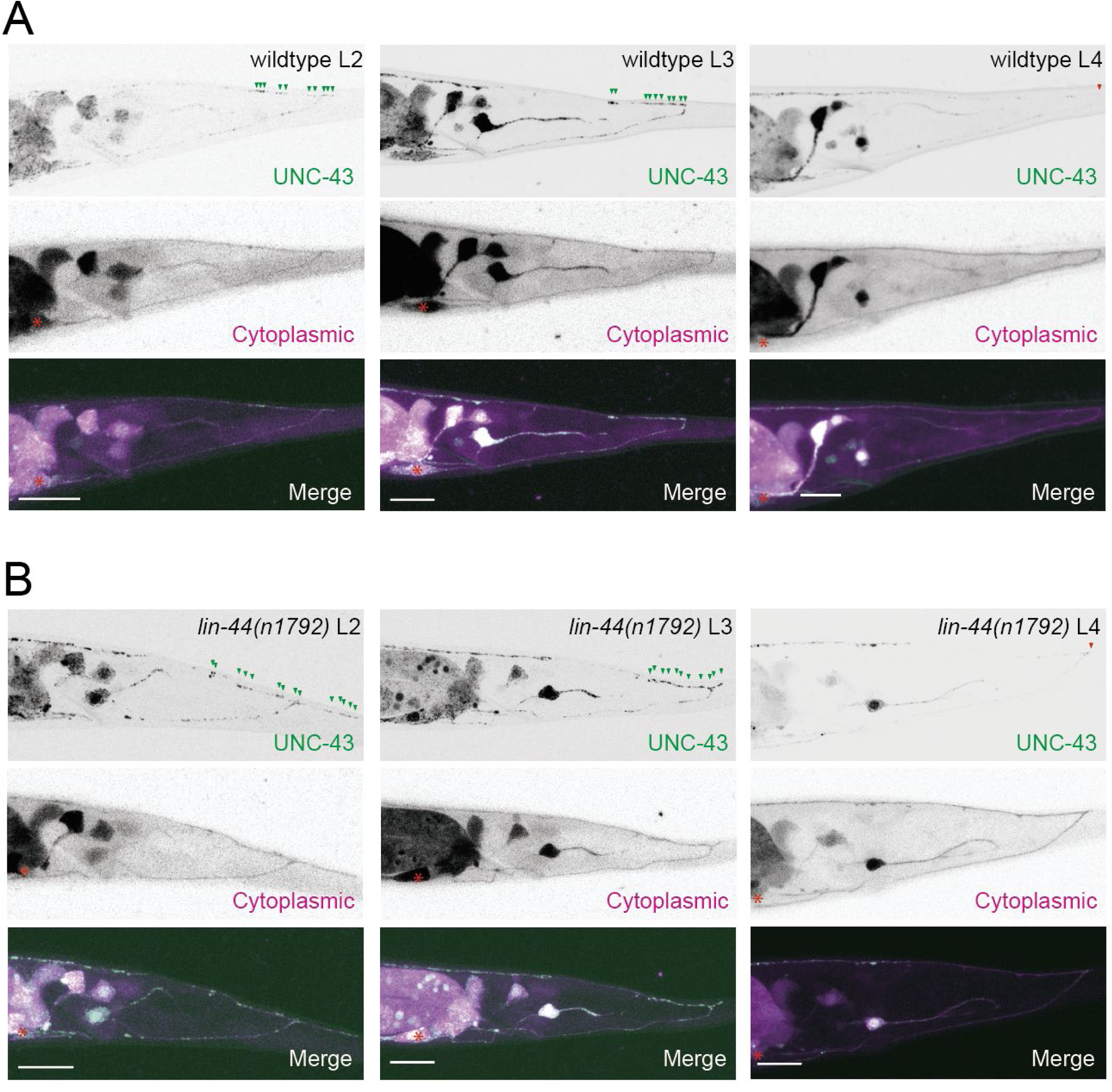
Endogenous UNC-43 is localized to the posterior region of PDB during neurite pruning. **(A, B)** Representative images of UNC-43 localization (top panels), PDB neurite structure labeled with cytoplasmic mScarlet (middle panels) and merged images (bottom panels) in wildtype **(A)** and *lin-44(n1792)* **(B)** animals at the L2 (left), L3(middle) and L4(right) stages. Green arrowheads indicate the UNC-43::7xGFP puncta around the posterior region of PDB neurites, and red arrowheads indicate the posterior end of PDB neurites lacking UNC-43::7xGFP puncta at the L4 stage. Asterisks denote the PDB cell body. Scale bars: 10 μm.

**Figure S2.**
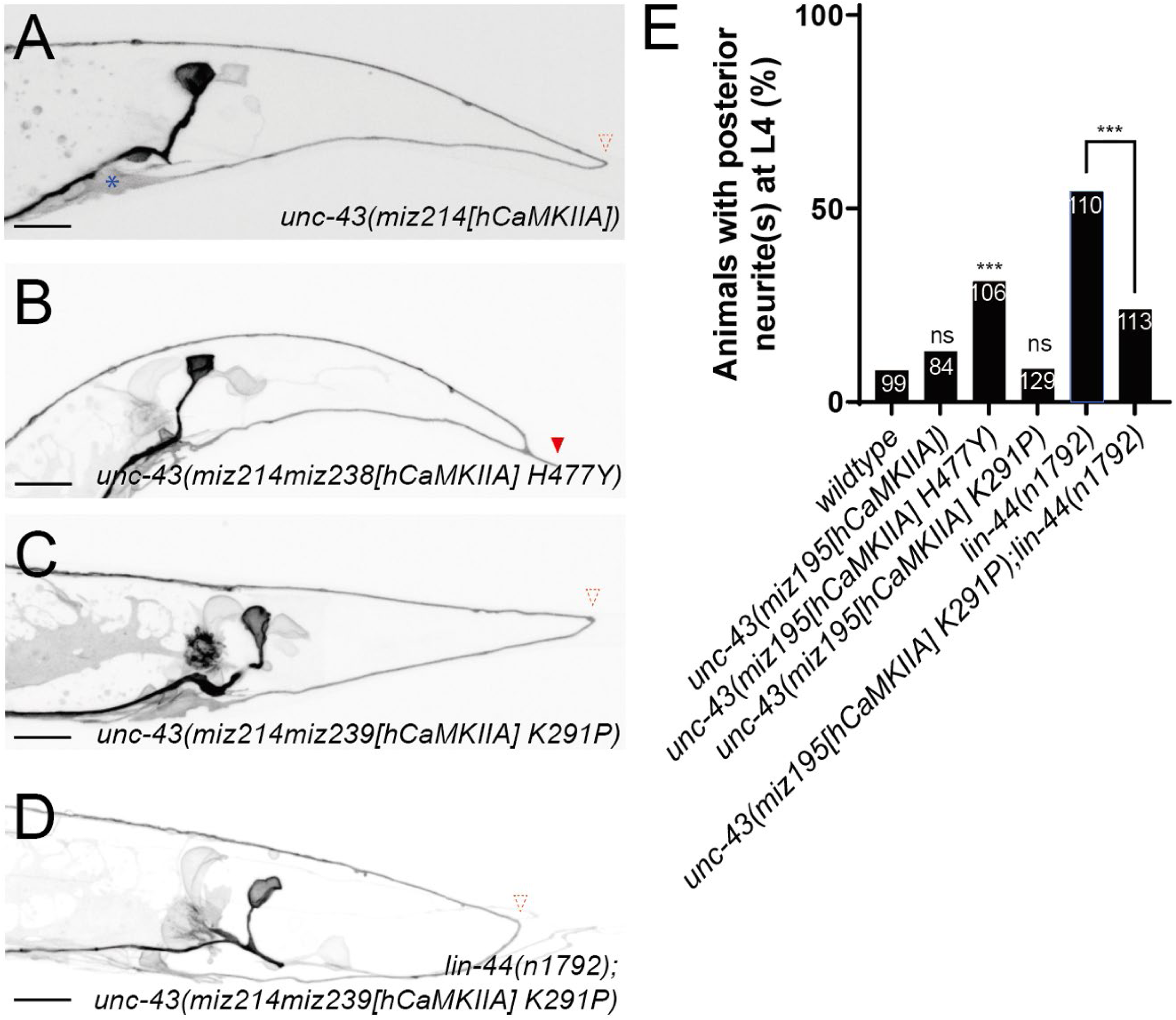
Human CaMKIIA functionally replaces *unc-43* in regulating PDB neurite pruning. **(A-D)** Representative images of PDB in *unc-43(miz214[hCaMKIIA])* (**A**)*, unc-43(miz214miz238[hCaMKIIA H477Y])* (**B**)*, unc-43(miz214miz209[hCaMKIIA K291P])* (**C**)*, lin-44(n1792); unc-43(miz214miz209[hCaMKIIA K291P])* (**D**) at the L4 stage. Red dotted arrowheads represent the absence of posterior neurites, and the red arrowhead represents the ectopic posterior neurite. Asterisks denote the PDB cell body. **(E)** Quantification of the animals with ectopic posterior neurites. Sample numbers are shown in each bar. ***p < 0.001; ns, not significant (Fisher’s exact test). Note the quantifications of wildtype and *lin-44(n1792)* are from Figure 1G. Scale bars: 10 μm.

**Figure S3.**
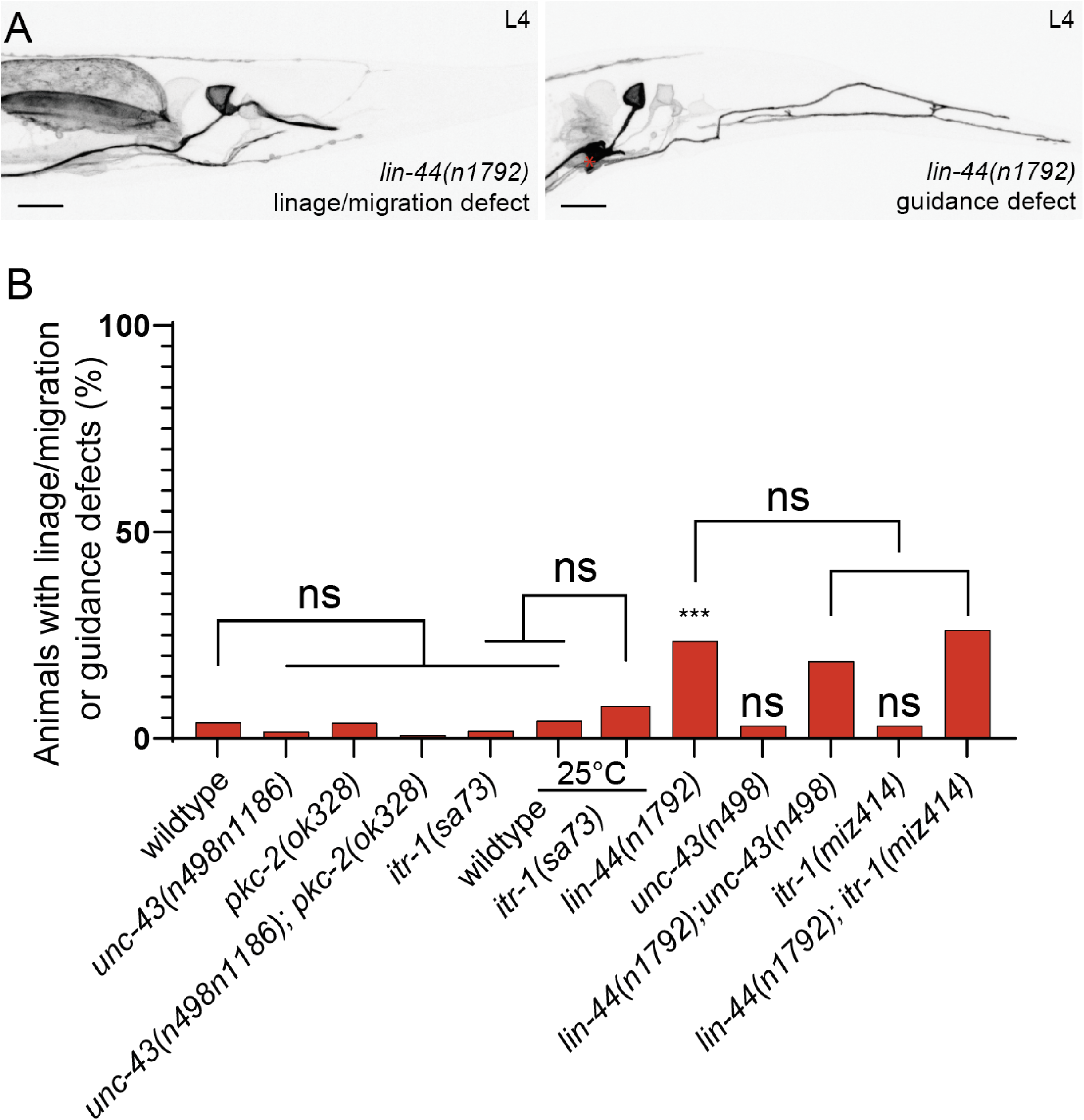
Wnt-calcium pathway components are not required for PDB cell fate specification and neurite guidance. **(A)** Representative images of PDB lineage of *lin-44(n1792)* mutant animals with no PDB neuron due to cell fate specification defect (left) and guidance defect (right). An asterisk represents the PDB cell body. **(B)** Quantification of the animals with lineage or guidance defects at the L4 stage. ***p < 0.001; ns, not significant (Fisher’s exact test). Scale bars: 10 μm.

**Figure S4.**
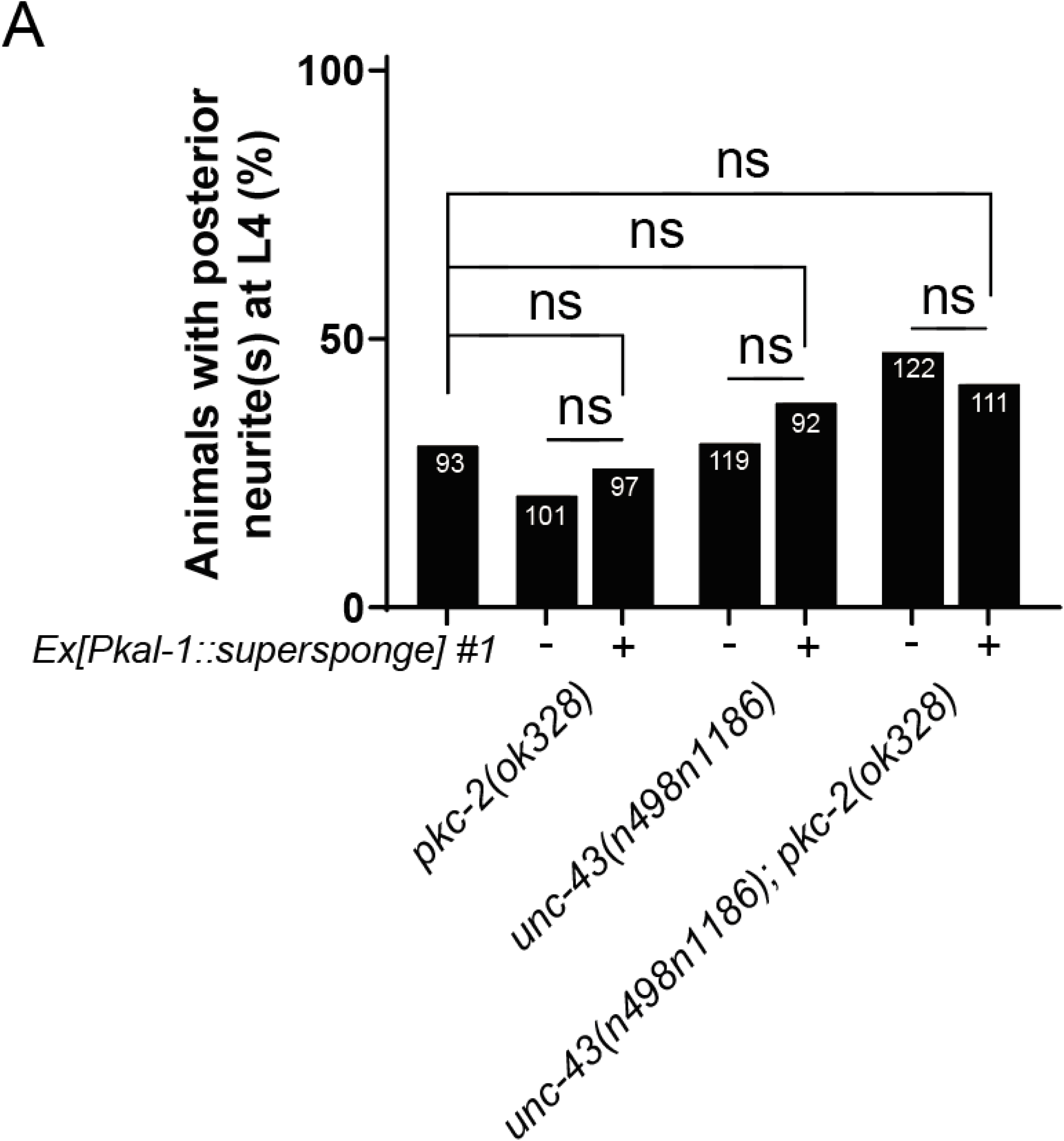
IP3 supersponge does not enhance the posterior neurite phenotypes of *unc-43* and *pkc-2*. Quantification of the animals with ectopic posterior neurites at the L4 stage. Sample numbers are shown in each bar. ns, not significant (Fisher’s exact test). Note the quantifications of *pkc-2(ok328)*, *unc-43(n498n1186)* and *pkc-2(ok328); unc-43(n498n1186)* are from Figure 1G.

**Figure S5.**
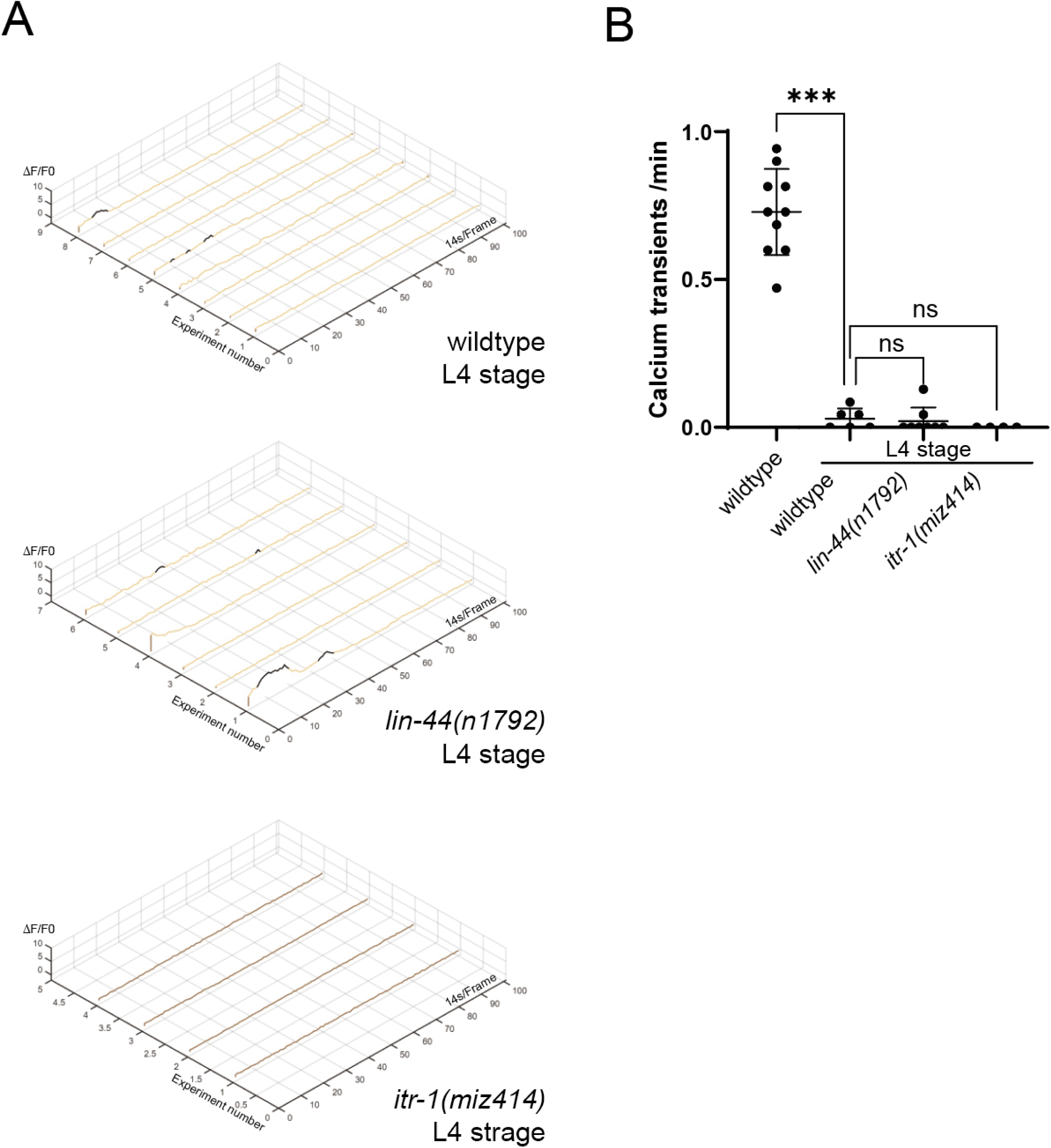
The frequency of calcium transients in PDB decreases at the L4 stage. **(A)** Change in GCaMP7s fluorescence within the ROI represented by ΔF/F0 in wildtype (top), *lin-44(n1792)* (middle), and *itr-1(miz414)* (bottom) animals at the L4 stage. Black segments of the plot denote the recognized calcium transients. **(B)** Plot of calcium transient frequencies. *** p < 0.001; ns, not significant (one-way ANOVA test).

**Figure S6.**
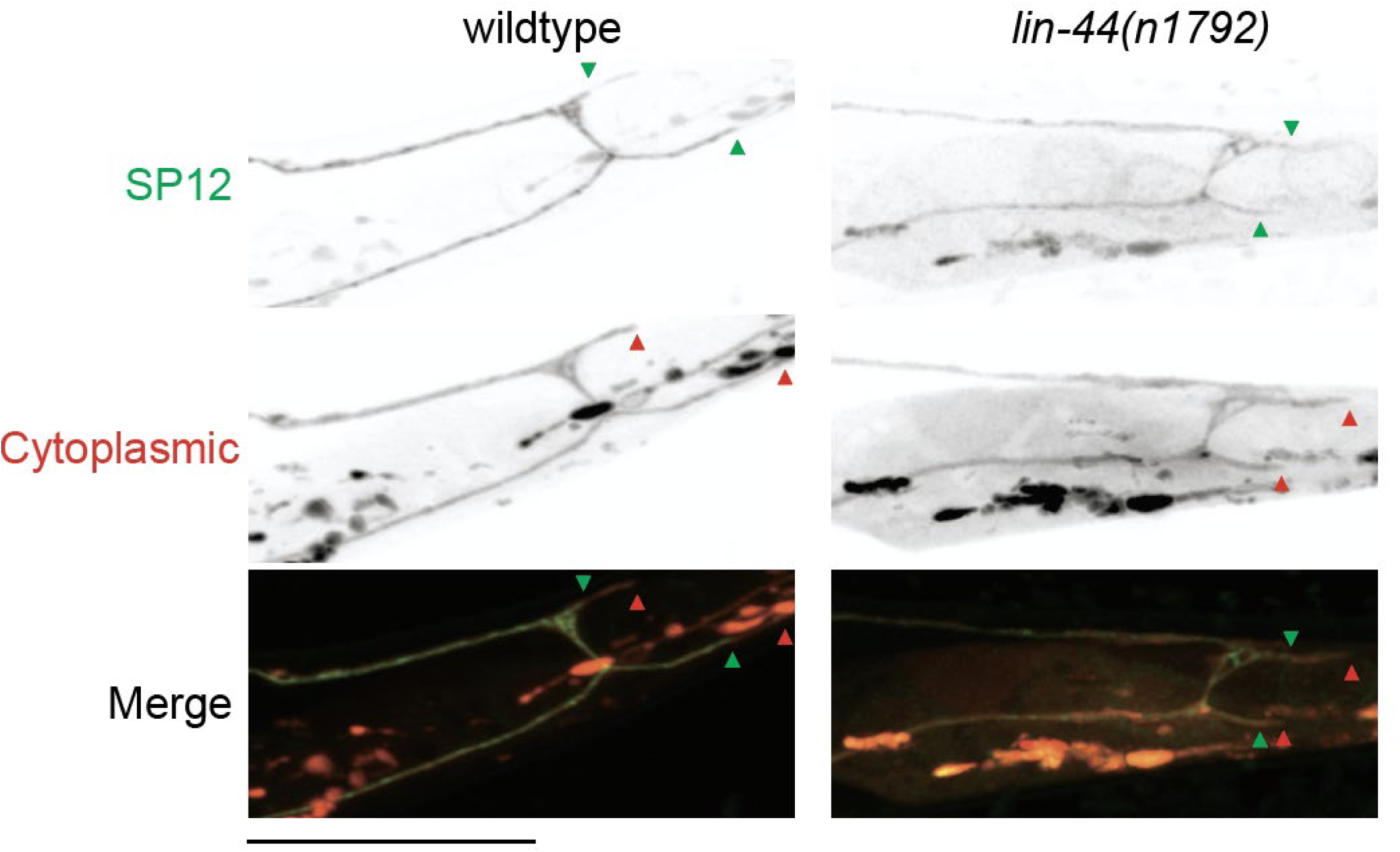
ER is localized at the posterior neurite in PDB during neurite pruning. **(A, B)** Representative images of GFPnovo2::SP12 (top panels), cytoplasmic mScarlet (middle panels), and merged images (bottom panels) in wild-type (**A**) and *lin-44(n1792)* (**B**) animals at the L2 stage. Green arrowheads indicate the posterior boundary of the SP12::GFPnovo2 signal, and red arrowheads denote the tips of posterior neurites of the PDB. Scale bars: 10 μm.

## Notes

### Competing Interest Statement

The authors have declared no competing interest.

## References

Antonini, A., & Stryker, M. P. (1993). Rapid remodeling of axonal arbors in the visual cortex. Science, 260(5115), 1819–1821. 10.1126/science.8511592

Artimovich, E., Jackson, R. K., Kilander, M. B. C., Lin, Y. C., & Nestor, M. W. (2017). PeakCaller: an automated graphical interface for the quantification of intracellular calcium obtained by high-content screening. BMC Neurosci, 18(1), 72. 10.1186/s12868-017-0391-y

Au, V., Li-Leger, E., Raymant, G., Flibotte, S., Chen, G., Martin, K., Fernando, L., Doell, C., Rosell, F. I., Wang, S., Edgley, M. L., Rougvie, A. E., Hutter, H., & Moerman, D. G. (2019). CRISPR/Cas9 Methodology for the Generation of Knockout Deletions in Caenorhabditis elegans. G3 (Bethesda), 9(1), 135–144. 10.1534/g3.118.200778

Bagri, A., Cheng, H. J., Yaron, A., Pleasure, S. J., & Tessier-Lavigne, M. (2003). Stereotyped pruning of long hippocampal axon branches triggered by retraction inducers of the semaphorin family. Cell, 113(3), 285–299. 10.1016/s0092-8674(03)00267-8

Baschieri, F., Porshneva, K., & Montagnac, G. (2020). Frustrated clathrin-mediated endocytosis - causes and possible functions. J Cell Sci, 133(11). 10.1242/jcs.240861

Baylis, H. A., Furuichi, T., Yoshikawa, F., Mikoshiba, K., & Sattelle, D. B. (1999). Inositol 1,4,5-trisphosphate receptors are strongly expressed in the nervous system, pharynx, intestine, gonad and excretory cell of Caenorhabditis elegans and are encoded by a single gene (itr-1). J Mol Biol, 294(2), 467–476. 10.1006/jmbi.1999.3229

Baylis, H. A., & Vazquez-Manrique, R. P. (2012). Genetic analysis of IP3 and calcium signalling pathways in C. elegans. Biochim Biophys Acta, 1820(8), 1253–1268. 10.1016/j.bbagen.2011.11.009

Bezprozvanny, I., Watras, J., & Ehrlich, B. E. (1991). Bell-shaped calcium-response curves of Ins(1,4,5)P3- and calcium-gated channels from endoplasmic reticulum of cerebellum. Nature, 351(6329), 751–754. 10.1038/351751a0

Bodakuntla, S., Nedozralova, H., Basnet, N., & Mizuno, N. (2021). Cytoskeleton and Membrane Organization at Axon Branches. Front Cell Dev Biol, 9, 707486. 10.3389/fcell.2021.707486

Brenner, S. (1974). The genetics of Caenorhabditis elegans. Genetics, 77(1), 71–94. 10.1093/genetics/77.1.71

Brunt, L., & Scholpp, S. (2018). The function of endocytosis in Wnt signaling. Cell Mol Life Sci, 75(5), 785–795. 10.1007/s00018-017-2654-2

Chia, P. H., Zhong, F. L., Niwa, S., Bonnard, C., Utami, K. H., Zeng, R., Lee, H., Eskin, A., Nelson, S. F., Xie, W. H., Al-Tawalbeh, S., El-Khateeb, M., Shboul, M., Pouladi, M. A., Al-Raqad, M., & Reversade, B. (2018). A homozygous loss-of-function CAMK2A mutation causes growth delay, frequent seizures and severe intellectual disability. Elife, 7. 10.7554/eLife.32451

Clandinin, T. R., DeModena, J. A., & Sternberg, P. W. (1998). Inositol trisphosphate mediates a RAS-independent response to LET-23 receptor tyrosine kinase activation in C. elegans. Cell, 92(4), 523–533. 10.1016/s0092-8674(00)80945-9

Coe, B. P., Stessman, H. A. F., Sulovari, A., Geisheker, M. R., Bakken, T. E., Lake, A. M., Dougherty, J. D., Lein, E. S., Hormozdiari, F., Bernier, R. A., & Eichler, E. E. (2019). Neurodevelopmental disease genes implicated by de novo mutation and copy number variation morbidity. Nat Genet, 51(1), 106–116. 10.1038/s41588-018-0288-4

Dana, H., Sun, Y., Mohar, B., Hulse, B. K., Kerlin, A. M., Hasseman, J. P., Tsegaye, G., Tsang, A., Wong, A., Patel, R., Macklin, J. J., Chen, Y., Konnerth, A., Jayaraman, V., Looger, L. L., Schreiter, E. R., Svoboda, K., & Kim, D. S. (2019). High-performance calcium sensors for imaging activity in neuronal populations and microcompartments. Nat Methods, 16(7), 649–657. 10.1038/s41592-019-0435-6

De Rubeis, S., He, X., Goldberg, A. P., Poultney, C. S., Samocha, K., Cicek, A. E., Kou, Y., Liu, L., Fromer, M., Walker, S., Singh, T., Klei, L., Kosmicki, J., Shih-Chen, F., Aleksic, B., Biscaldi, M., Bolton, P. F., Brownfeld, J. M., Cai, J.,…Buxbaum, J. D. (2014). Synaptic, transcriptional and chromatin genes disrupted in autism. Nature, 515(7526), 209–215. 10.1038/nature13772

Delos Santos, R. C., Bautista, S., Lucarelli, S., Bone, L. N., Dayam, R. M., Abousawan, J., Botelho, R. J., & Antonescu, C. N. (2017). Selective regulation of clathrin-mediated epidermal growth factor receptor signaling and endocytosis by phospholipase C and calcium. Mol Biol Cell, 28(21), 2802–2818. 10.1091/mbc.E16-12-0871

Dhavan, R., Greer, P. L., Morabito, M. A., Orlando, L. R., & Tsai, L. H. (2002). The cyclin-dependent kinase 5 activators p35 and p39 interact with the alpha-subunit of Ca2+/calmodulin-dependent protein kinase II and alpha-actinin-1 in a calcium-dependent manner. J Neurosci, 22(18), 7879–7891. 10.1523/JNEUROSCI.22-18-07879.2002

Fire, A. (1986). Integrative transformation of Caenorhabditis elegans. EMBO J, 5(10), 2673–2680. 10.1002/j.1460-2075.1986.tb04550.x

Ghanta, K. S., & Mello, C. C. (2020). Melting dsDNA Donor Molecules Greatly Improves Precision Genome Editing in Caenorhabditis elegans. Genetics, 216(3), 643–650. 10.1534/genetics.120.303564

Ghosh-Roy, A., Wu, Z., Goncharov, A., Jin, Y., & Chisholm, A. D. (2010). Calcium and cyclic AMP promote axonal regeneration in Caenorhabditis elegans and require DLK-1 kinase. J Neurosci, 30(9), 3175–3183. 10.1523/JNEUROSCI.5464-09.2010

Hayashi, Y., Hirotsu, T., Iwata, R., Kage-Nakadai, E., Kunitomo, H., Ishihara, T., Iino, Y., & Kubo, T. (2009). A trophic role for Wnt-Ror kinase signaling during developmental pruning in Caenorhabditis elegans. Nat Neurosci, 12(8), 981–987. 10.1038/nn.2347

He, C. W., Liao, C. P., & Pan, C. L. (2018). Wnt signalling in the development of axon, dendrites and synapses. Open Biol, 8(10). 10.1098/rsob.180116

He, S., Cuentas-Condori, A., & Miller, D. M., 3rd. (2019). NATF (Native and Tissue-Specific Fluorescence): A Strategy for Bright, Tissue-Specific GFP Labeling of Native Proteins in Caenorhabditis elegans. Genetics, 212(2), 387–395. 10.1534/genetics.119.302063

Heisenberg, C. P., Tada, M., Rauch, G. J., Saude, L., Concha, M. L., Geisler, R., Stemple, D. L., Smith, J. C., & Wilson, S. W. (2000). Silberblick/Wnt11 mediates convergent extension movements during zebrafish gastrulation. Nature, 405(6782), 76–81. 10.1038/35011068

Helbig, I., Lopez-Hernandez, T., Shor, O., Galer, P., Ganesan, S., Pendziwiat, M., Rademacher, A., Ellis, C. A., Humpfer, N., Schwarz, N., Seiffert, S., Peeden, J., Shen, J., Sterbova, K., Hammer, T. B., Moller, R. S., Shinde, D. N., Tang, S., Smith, L.,…Consortium, G. (2019). A Recurrent Missense Variant in AP2M1 Impairs Clathrin-Mediated Endocytosis and Causes Developmental and Epileptic Encephalopathy. Am J Hum Genet, 104(6), 1060–1072. 10.1016/j.ajhg.2019.04.001

Hensch, T. K. (2005). Critical period plasticity in local cortical circuits. Nat Rev Neurosci, 6(11), 877–888. 10.1038/nrn1787

Hutchins, B. I., Li, L., & Kalil, K. (2011). Wnt/calcium signaling mediates axon growth and guidance in the developing corpus callosum. Dev Neurobiol, 71(4), 269–283. 10.1002/dneu.20846

Jiang, L. I., & Sternberg, P. W. (1998). Interactions of EGF, Wnt and HOM-C genes specify the P12 neuroectoblast fate in C. elegans. Development, 125(12), 2337–2347. 10.1242/dev.125.12.2337

Kage, E., Hayashi, Y., Takeuchi, H., Hirotsu, T., Kunitomo, H., Inoue, T., Arai, H., Iino, Y., & Kubo, T. (2005). MBR-1, a novel helix-turn-helix transcription factor, is required for pruning excessive neurites in Caenorhabditis elegans. Curr Biol, 15(17), 1554–1559. 10.1016/j.cub.2005.07.057

Koushika, S. P., & Nonet, M. L. (2000). Sorting and transport in C. elegans: aA model system with a sequenced genome. Curr Opin Cell Biol, 12(4), 517–523. 10.1016/s0955-0674(00)00125-3

Kuhl, M., Sheldahl, L. C., Malbon, C. C., & Moon, R. T. (2000). Ca(2+)/calmodulin-dependent protein kinase II is stimulated by Wnt and Frizzled homologs and promotes ventral cell fates in Xenopus. J Biol Chem, 275(17), 12701–12711. 10.1074/jbc.275.17.12701

Kuhl, M., Sheldahl, L. C., Park, M., Miller, J. R., & Moon, R. T. (2000). The Wnt/Ca2+ pathway: a new vertebrate Wnt signaling pathway takes shape. Trends Genet, 16(7), 279–283. 10.1016/s0168-9525(00)02028-x

Kuo, C. T., Jan, L. Y., & Jan, Y. N. (2005). Dendrite-specific remodeling of Drosophila sensory neurons requires matrix metalloproteases, ubiquitin-proteasome, and ecdysone signaling. Proc Natl Acad Sci U S A, 102(42), 15230–15235. 10.1073/pnas.0507393102

Kurashina, M., & Mizumoto, K. (2023). Targeting endogenous proteins for spatial and temporal knockdown using auxin-inducible degron in Caenorhabditis elegans. STAR Protoc, 4(1), 102028. 10.1016/j.xpro.2022.102028

Kurashina, M., Snow, A. W., & Mizumoto, K. (2025). A modular system to label endogenous presynaptic proteins using split fluorophores in Caenorhabditis elegans. Genetics, 229(3). 10.1093/genetics/iyae214

Kury, S., van Woerden, G. M., Besnard, T., Proietti Onori, M., Latypova, X., Towne, M. C., Cho, M. T., Prescott, T. E., Ploeg, M. A., Sanders, S., Stessman, H. A. F., Pujol, A., Distel, B., Robak, L. A., Bernstein, J. A., Denomme-Pichon, A. S., Lesca, G., Sellars, E. A., Berg, J.,…Mercier, S. (2017). De Novo Mutations in Protein Kinase Genes CAMK2A and CAMK2B Cause Intellectual Disability. Am J Hum Genet, 101(5), 768–788. 10.1016/j.ajhg.2017.10.003

LeBoeuf, B., Chen, X., & Garcia, L. R. (2020). WNT regulates programmed muscle remodeling through PLC-beta and calcineurin in Caenorhabditis elegans males. Development, 147(9). 10.1242/dev.181305

Lee, J., Jongeward, G. D., & Sternberg, P. W. (1994). unc-101, a gene required for many aspects of Caenorhabditis elegans development and behavior, encodes a clathrin-associated protein. Genes Dev, 8(1), 60–73. 10.1101/gad.8.1.60

Lee, R. Y., Lobel, L., Hengartner, M., Horvitz, H. R., & Avery, L. (1997). Mutations in the alpha1 subunit of an L-type voltage-activated Ca2+ channel cause myotonia in Caenorhabditis elegans. EMBO J, 16(20), 6066–6076. 10.1093/emboj/16.20.6066

Lehmann, K. S., Hupp, M. T., Abalde-Atristain, L., Jefferson, A., Cheng, Y. C., Sheehan, A. E., Kang, Y., & Freeman, M. R. (2025). Astrocyte-dependent local neurite pruning in Beat-Va neurons. J Cell Biol, 224(1). 10.1083/jcb.202312043

Li, L., Hutchins, B. I., & Kalil, K. (2009). Wnt5a induces simultaneous cortical axon outgrowth and repulsive axon guidance through distinct signaling mechanisms. J Neurosci, 29(18), 5873–5883. 10.1523/JNEUROSCI.0183-09.2009

Liu, X., Guo, X., Niu, L., Li, X., Sun, F., Hu, J., Wang, X., & Shen, K. (2019). Atlastin-1 regulates morphology and function of endoplasmic reticulum in dendrites. Nat Commun, 10(1), 568. 10.1038/s41467-019-08478-6

Lu, M., & Mizumoto, K. (2019). Gradient-independent Wnt signaling instructs asymmetric neurite pruning in C. elegans. Elife, 8. 10.7554/eLife.50583

Luo, L., & O’Leary, D. D. (2005). Axon retraction and degeneration in development and disease. Annu Rev Neurosci, 28, 127–156. 10.1146/annurev.neuro.28.061604.135632

Malaiwong, N., Porta-de-la-Riva, M., & Krieg, M. (2023). FLInt: single shot safe harbor transgene integration via Fluorescent Landmark Interference. G3 (Bethesda), 13(5). 10.1093/g3journal/jkad041

Mariol, M. C., Walter, L., Bellemin, S., & Gieseler, K. (2013). A rapid protocol for integrating extrachromosomal arrays with high transmission rate into the C. elegans genome. J Vis Exp(82), e50773. 10.3791/50773

Maryon, E. B., Coronado, R., & Anderson, P. (1996). unc-68 encodes a ryanodine receptor involved in regulating C. elegans body-wall muscle contraction. J Cell Biol, 134(4), 885–893. 10.1083/jcb.134.4.885

Mayseless, O., Berns, D. S., Yu, X. M., Riemensperger, T., Fiala, A., & Schuldiner, O. (2018). Developmental Coordination during Olfactory Circuit Remodeling in Drosophila. Neuron, 99(6), 1204–1215 e1205. 10.1016/j.neuron.2018.07.050

Mayseless, O., Shapira, G., Rachad, E. Y., Fiala, A., & Schuldiner, O. (2023). Neuronal excitability as a regulator of circuit remodeling. Curr Biol, 33(5), 981–989 e983. 10.1016/j.cub.2023.01.032

McQuate, A., Latorre-Esteves, E., & Barria, A. (2017). A Wnt/Calcium Signaling Cascade Regulates Neuronal Excitability and Trafficking of NMDARs. Cell Rep, 21(1), 60–69. 10.1016/j.celrep.2017.09.023

McVicker, D. P., Millette, M. M., & Dent, E. W. (2015). Signaling to the microtubule cytoskeleton: an unconventional role for CaMKII. Dev Neurobiol, 75(4), 423–434. 10.1002/dneu.22227

Mello, C. C., Kramer, J. M., Stinchcomb, D., & Ambros, V. (1991). Efficient gene transfer in C.elegans: extrachromosomal maintenance and integration of transforming sequences. EMBO J, 10(12), 3959–3970. 10.1002/j.1460-2075.1991.tb04966.x

Pan, M., Liu, P. W., Ozawa, Y., Arima-Yoshida, F., Dong, G., Sawahata, M., Mori, D., Nagase, M., Fujii, H., Ueda, S., Yabuuchi, Y., Liu, X., Narita, H., Konno, A., Hirai, H., Ozaki, N., Yamada, K., Kidokoro, H., Bito, H.,…Takemoto-Kimura, S. (2025). A hyper-activatable CAMK2A variant associated with intellectual disability causes exaggerated long-term potentiation and learning impairments. Transl Psychiatry, 15(1), 95. 10.1038/s41398-025-03316-4

Park, M., & Shen, K. (2012). WNTs in synapse formation and neuronal circuitry. EMBO J, 31(12), 2697–2704. 10.1038/emboj.2012.145

Peng, Y., Lee, J., Rowland, K., Wen, Y., Hua, H., Carlson, N., Lavania, S., Parrish, J. Z., & Kim, M. D. (2015). Regulation of dendrite growth and maintenance by exocytosis. J Cell Sci, 128(23), 4279–4292. 10.1242/jcs.174771

Penzes, P., Cahill, M. E., Jones, K. A., VanLeeuwen, J. E., & Woolfrey, K. M. (2011). Dendritic spine pathology in neuropsychiatric disorders. Nat Neurosci, 14(3), 285–293. 10.1038/nn.2741

Proietti Onori, M., Koopal, B., Everman, D. B., Worthington, J. D., Jones, J. R., Ploeg, M. A., Mientjes, E., van Bon, B. W., Kleefstra, T., Schulman, H., Kushner, S. A., Kury, S., Elgersma, Y., & van Woerden, G. M. (2018). The intellectual disability-associated CAMK2G p.Arg292Pro mutation acts as a pathogenic gain-of-function. Hum Mutat, 39(12), 2008–2024. 10.1002/humu.23647

Reiner, D. J., Newton, E. M., Tian, H., & Thomas, J. H. (1999). Diverse behavioural defects caused by mutations in Caenorhabditis elegans unc-43 CaM kinase II. Nature, 402(6758), 199–203. 10.1038/46072

Riccomagno, M. M., & Kolodkin, A. L. (2015). Sculpting neural circuits by axon and dendrite pruning. Annu Rev Cell Dev Biol, 31, 779–805. 10.1146/annurev-cellbio-100913-013038

Rigter, P. M. F., de Konink, C., Dunn, M. J., Proietti Onori, M., Humberson, J. B., Thomas, M., Barnes, C., Prada, C. E., Weaver, K. N., Ryan, T. D., Caluseriu, O., Conway, J., Calamaro, E., Fong, C. T., Wuyts, W., Meuwissen, M., Hordijk, E., Jonkers, C. N., Anderson, L.,…van Woerden, G. M. (2024). Role of CAMK2D in neurodevelopment and associated conditions. Am J Hum Genet, 111(2), 364–382. 10.1016/j.ajhg.2023.12.016

Robatzek, M., & Thomas, J. H. (2000). Calcium/calmodulin-dependent protein kinase II regulates Caenorhabditis elegans locomotion in concert with a G(o)/G(q) signaling network. Genetics, 156(3), 1069–1082. 10.1093/genetics/156.3.1069

Salinas, P. C., & Zou, Y. (2008). Wnt signaling in neural circuit assembly. Annu Rev Neurosci, 31, 339–358. 10.1146/annurev.neuro.31.060407.125649

Saneyoshi, T., Kume, S., Amasaki, Y., & Mikoshiba, K. (2002). The Wnt/calcium pathway activates NF-AT and promotes ventral cell fate in Xenopus embryos. Nature, 417(6886), 295–299. 10.1038/417295a

Sato, M., Sato, K., Fonarev, P., Huang, C. J., Liou, W., & Grant, B. D. (2005). Caenorhabditis elegans RME-6 is a novel regulator of RAB-5 at the clathrin-coated pit. Nat Cell Biol, 7(6), 559–569. 10.1038/ncb1261

Schafer, W. R., & Kenyon, C. J. (1995). A calcium-channel homologue required for adaptation to dopamine and serotonin in Caenorhabditis elegans. Nature, 375(6526), 73–78. 10.1038/375073a0

Schafer, W. R., Sanchez, B. M., & Kenyon, C. J. (1996). Genes affecting sensitivity to serotonin in Caenorhabditis elegans. Genetics, 143(3), 1219–1230. 10.1093/genetics/143.3.1219

Schuldiner, O., & Yaron, A. (2015). Mechanisms of developmental neurite pruning. Cell Mol Life Sci, 72(1), 101–119. 10.1007/s00018-014-1729-6

Sheldahl, L. C., Park, M., Malbon, C. C., & Moon, R. T. (1999). Protein kinase C is differentially stimulated by Wnt and Frizzled homologs in a G-protein-dependent manner. Curr Biol, 9(13), 695–698. 10.1016/s0960-9822(99)80310-8

Sheldahl, L. C., Slusarski, D. C., Pandur, P., Miller, J. R., Kuhl, M., & Moon, R. T. (2003). Dishevelled activates Ca2+ flux, PKC, and CamKII in vertebrate embryos. J Cell Biol, 161(4), 769–777. 10.1083/jcb.200211094

Shim, J., & Lee, J. (2005). The AP-3 Clathrin-associated Complex Is Essential for Embryonic and Larval Development in Caenorhabditis elegans. Molecules and Cells, 19(3), 452–457. 10.1016/s1016-8478(23)13192-x

Slusarski, D. C., Corces, V. G., & Moon, R. T. (1997). Interaction of Wnt and a Frizzled homologue triggers G-protein-linked phosphatidylinositol signalling. Nature, 390(6658), 410–413. 10.1038/37138

Slusarski, D. C., Yang-Snyder, J., Busa, W. B., & Moon, R. T. (1997). Modulation of embryonic intracellular Ca2+ signaling by Wnt-5A. Dev Biol, 182(1), 114–120. 10.1006/dbio.1996.8463

Stankovic, I., Smit, P., Cross, J., Rai, A., Wolujewicz, P., Greening, D., & Colak, D. (2025). Extracellular vesicle profiling reveals novel autism signatures in patient-derived forebrain organoids. Transl Psychiatry, 15(1), 393. 10.1038/s41398-025-03607-w

Steger, K. A., Shtonda, B. B., Thacker, C., Snutch, T. P., & Avery, L. (2005). The C. elegans T-type calcium channel CCA-1 boosts neuromuscular transmission. J Exp Biol, 208(Pt 11), 2191–2203. 10.1242/jeb.01616

Tojima, T., Itofusa, R., & Kamiguchi, H. (2010). Asymmetric clathrin-mediated endocytosis drives repulsive growth cone guidance. Neuron, 66(3), 370–377. 10.1016/j.neuron.2010.04.007

Tojima, T., Itofusa, R., & Kamiguchi, H. (2014). Steering neuronal growth cones by shifting the imbalance between exocytosis and endocytosis. J Neurosci, 34(21), 7165–7178. 10.1523/JNEUROSCI.5261-13.2014

Tojima, T., & Kamiguchi, H. (2015). Exocytic and endocytic membrane trafficking in axon development. Dev Growth Differ, 57(4), 291–304. 10.1111/dgd.12218

Walker, D. S., Gower, N. J., Ly, S., Bradley, G. L., & Baylis, H. A. (2002). Regulated disruption of inositol 1,4,5-trisphosphate signaling in Caenorhabditis elegans reveals new functions in feeding and embryogenesis. Mol Biol Cell, 13(4), 1329–1337. 10.1091/mbc.01-08-0422

Walsh, M. K., & Lichtman, J. W. (2003). In vivo time-lapse imaging of synaptic takeover associated with naturally occurring synapse elimination. Neuron, 37(1), 67–73. 10.1016/s0896-6273(02)01142-x

Westfall, T. A., Brimeyer, R., Twedt, J., Gladon, J., Olberding, A., Furutani-Seiki, M., & Slusarski, D. C. (2003). Wnt-5/pipetail functions in vertebrate axis formation as a negative regulator of Wnt/beta-catenin activity. J Cell Biol, 162(5), 889–898. 10.1083/jcb.200303107

White, J. G., Southgate, E., Thomson, J. N., & Brenner, S. (1986). The structure of the nervous system of the nematode Caenorhabditis elegans. Philos Trans R Soc Lond B Biol Sci, 314(1165), 1–340. 10.1098/rstb.1986.0056

Xu, N. J., & Henkemeyer, M. (2009). Ephrin-B3 reverse signaling through Grb4 and cytoskeletal regulators mediates axon pruning. Nat Neurosci, 12(3), 268–276. 10.1038/nn.2254

Yamahachi, H., Marik, S. A., McManus, J. N., Denk, W., & Gilbert, C. D. (2009). Rapid axonal sprouting and pruning accompany functional reorganization in primary visual cortex. Neuron, 64(5), 719–729. 10.1016/j.neuron.2009.11.026

Yamazaki, H., Koganezawa, N., Yokoo, H., Sekino, Y., & Shirao, T. (2024). Super-resolution imaging reveals the relationship between CaMKIIbeta and drebrin within dendritic spines. Neurosci Res, 199, 30–35. 10.1016/j.neures.2023.08.002

Zhang, H., Lesnov, G. D., Subach, O. M., Zhang, W., Kuzmicheva, T. P., Vlaskina, A. V., Samygina, V. R., Chen, L., Ye, X., Nikolaeva, A. Y., Gabdulkhakov, A., Papadaki, S., Qin, W., Borshchevskiy, V., Perfilov, M. M., Gavrikov, A. S., Drobizhev, M., Mishin, A. S., Piatkevich, K. D., & Subach, F. V. (2024). Bright and stable monomeric green fluorescent protein derived from StayGold. Nat Methods, 21(4), 657–665. 10.1038/s41592-024-02203-y

Zhou, Y., Lai, B., & Gan, W. B. (2017). Monocular deprivation induces dendritic spine elimination in the developing mouse visual cortex. Sci Rep, 7(1), 4977. 10.1038/s41598-017-05337-6

